# Generalizable Protein Dynamics in Kinases: Physics is the key

**DOI:** 10.1101/2025.03.06.641878

**Authors:** Soumendranath Bhakat, Shray Vats, Andreas Mardt, Alexei Degterev

## Abstract

Kinases play crucial roles in signaling pathways across oncology, inflammation, and neurodegenerative diseases. Historically, their conformational states have been defined by the DFG motif: DFG_in_ and DFG_out_. However, this binary paradigm overlooks the broader conformational heterogeneity of apo kinases, which encompasses multiple metastable states within an expanded DFG-Phe ensemble. We introduced a novel nomenclature that integrates the dynamics of the DFG-Phe, activation loop, and *αC*-helix, highlighting how these regions respond to various ensemble perturbations (e.g., mutations, ligand binding, and protein–protein interactions). A major bottleneck in studying kinases lies in sampling their wide-ranging conformations because static snapshots often remain trapped in specific free energy minima, limiting traditional molecular dynamics simulations from exploring multiple functionally relevant states starting from a single structure. To accelerate conformational sampling, we introduce a computational framework that integrates AlphaFold, machine learning, physics-based simulations, and Markov state modeling. Rather than focusing on single-structure snapshots or structural hypotheses generated by protein structure prediction models, our framework captures shifts in conformational populations under varying perturbations, shedding light on both the thermodynamics and kinetics of the transitions. We show generalizability of our ensemble definitions and protocol across members of serine-threonine kinases and tyrosine kinases. A key innovation lies in our machine learning algorithms, which capture slowly varying structural features to uncover hidden states and generate latent representations of conformational motions across different kinase domains. The physics-refined structural ensemble sampled from the latent layers is then used to launch new simulations that more comprehensively explore the full conformational landscape than traditional molecular simulation approaches. By capturing how these conformational shifts influence downstream protein–protein interactions, conformational allostery, and cryptic pocket formation, our accelerated simulation framework provides deeper insights into generalized molecular recognition mechanism in kinases and how conformational heterogeneity is influenced by ensemble perturbations. This framework can be extended to investigate broader protein families—such as G protein-coupled receptors (GPCRs) and tumor necrosis factors (TNFs)—where functional outcomes are dictated by conformational heterogeneity.

## Introduction

Protein kinases regulate signaling by transferring ATP’s phosphate to target proteins. Humans encode over 500 serine/threonine kinases (STKs) that phosphorylate serine/threonine residues and engage in kinase-mediated protein-protein interactions controlling cell-cycle, metabolic, and stress pathways implicated in cancer, inflammation, and neurodegeneration.^1^. The activity of STKs is heavily modulated by their conformational dynamics, which are influenced by perturbations such as mutations, small molecule binding, and protein–protein interactions^2,3^. Tyrosine kinases (TKs) similarly phosphorylate tyrosine to drive growth-factor and immune signaling, and dysregulation of either class leads to several human disease, making both STKs and TKs prime therapeutic targets. The blockbuster success of the TK inhibitor imatinib (Gleevec) steered most computational work toward TK dynamics, leaving STKs under-explored.

Like most kinases, STKs contain a conserved Asp-Phe-Gly (DFG) motif essential for their function. Historically, kinase conformational dynamics have been described by the DFG_in_ and DFG_out_ states^4,5^, corresponding to active and inactive conformations, respectively. In the active (DFG_in_) conformation, the DFG-Phe residue is packed into a hydrophobic pocket, while the DFG-Asp faces outward. Conversely, in the inactive (DFG_out_) state, the DFG-Asp flips, and the DFG-Phe relocates out of the hydrophobic pocket. For STKs, the transition from DFG_in_ to DFG_out_ requires overcoming a significantly high free energy barrier and is often induced by the binding of type II kinase inhibitors^6,7^.

However, this binary classification overlooks the conformational heterogeneity of apo STKs, which predominantly exist in a dynamic equilibrium among multiple metastable states within a global DFG_in_ ensemble. The current nomenclature paradigm is inadequate to capture this conformational diversity. Understanding the populations and timescales associated with the conformational heterogeneity is essential for elucidating the effects of perturbations on STK dynamics, providing molecular-level insights into disease mechanisms, and facilitating the development of selective therapeutics targeting these kinases. Here, we propose a new nomenclature to define the conformational heterogeneity of STKs, which not only accounts for the conformational dynamics of the DFG-Phe residue but also encompasses the variability of other key structural motifs, namely the αC-helix and the activation loop, within the broader DFG-Phe ensemble.

A key challenge in sampling the conformational heterogeneity of apo STKs is that the timescales associated with transitions between metastable states are often too long to be captured by classical unbiased molecular dynamics (MD) simulations^8,9^. These transitions require navigating a rugged free energy landscape with numerous microstates, causing classical MD simulations to become trapped in local minima^10^. This issue is further exacerbated when studying how ligand binding to specific protein states modulates shift in populations and kinetics among functionally relevant metastable states (**Figure 1**). Traditional approaches are limited to specialized MD simulation hardware, such as Anton^11^, and struggle to capture the slowly varying structural features of different protein domains that govern conformational shifts in biomolecules.

**Figure 1.**
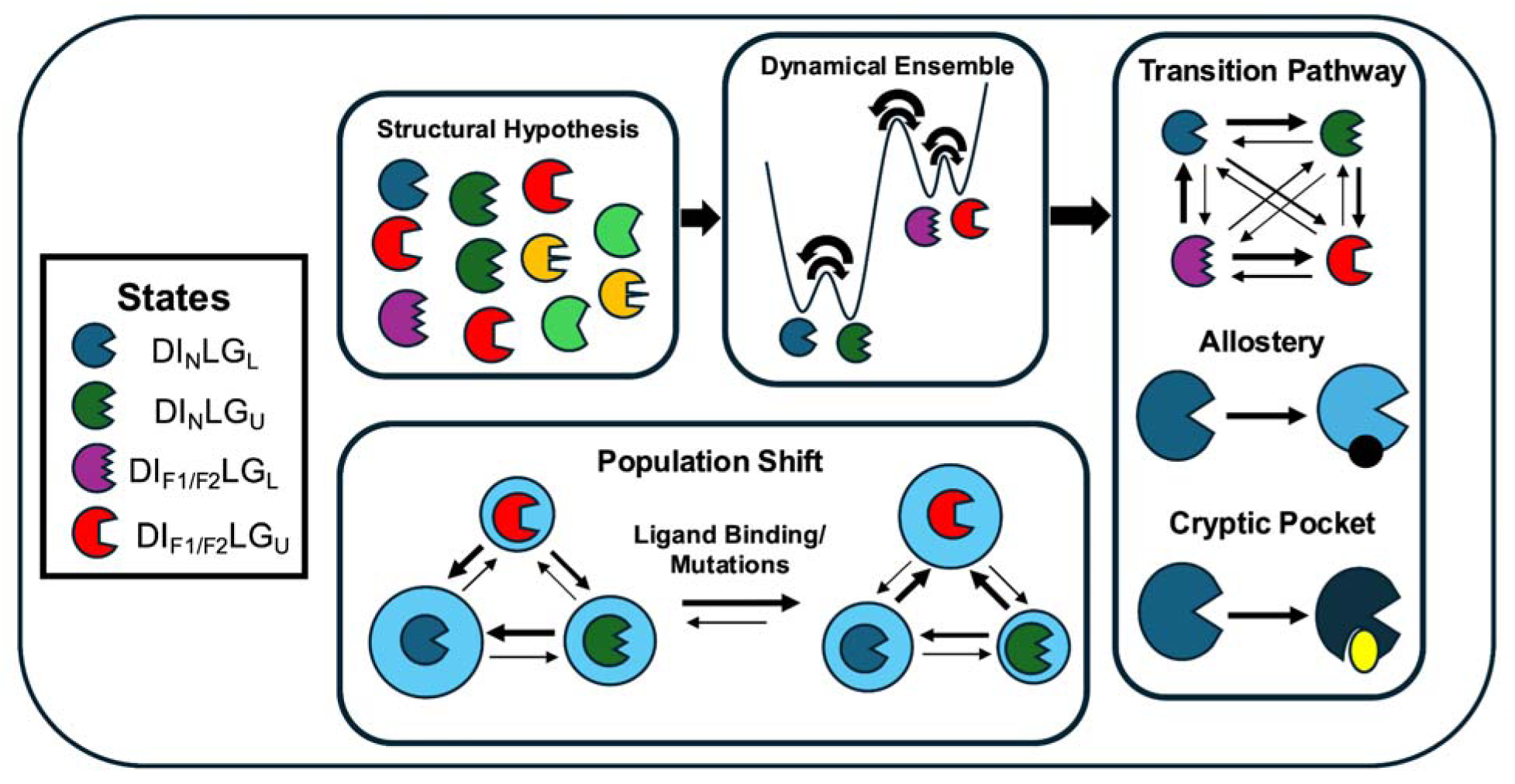
Our approach introduces an ensemble definition (states) of kinases, showing how AI/ML-augmented molecular simulations capture the conformational heterogeneity of kinase states and th effects of perturbations—such as mutations and ligand binding—on their populations and kinetics. While protein structure prediction models such as AlphaFold provide structural hypotheses, they lack th physics needed to accurately capture conformational dynamics. In contrast, physics-based molecular simulations informed by these structural hypotheses combined with machine learning reveal state populations, transition pathways, conformational allostery, and novel druggable cryptic pockets, expanding the scope of early-stage drug discovery.

To address this sampling challenge, we utilized a previously developed framework^12^ that employs multiple sequence alignment (MSA) subsampling^13,14^ with AlphaFold to generate conformational heterogeneity. This approach uses multiple independent MD simulations from the AlphaFold generated conformational ensemble and Markov state model (MSM)^15–17^ to sample the complete spectrum of conformational heterogeneity in apo STKs. However, this method can still become trapped in free energy basins if the initial structural diversity and subsequent MD simulations fail to overcome local free energy barriers.

To overcome this limitation, we developed a new adaptive sampling protocol^18^ based on training a time-lagged autoencoder on short, physics-based molecular simulations seeded from AlphaFold generated ensembles. This protocol captures a physics-refined diverse conformational ensemble, enabling the sampling of protein conformations from underexplored regions of conformational space. Molecular simulations starting from this physics-refined conformational ensemble allowed us to capture a complete conformational spectrum of different functionally relevant metastable states in STKs. The weights from the *time-lagged* autoencoder^19^, mapped to protein structures, enabled us to identify structurally important conformational features that can guide the interpretation of experimental results and inform the design of new experiments. Further, the ability to capture physics refined conformational ensemble rather than single structure allowed us to capture conformational allostery^8,20,21^ and cryptic pockets^12,22^ in previously undruggable kinase-protein complexes.

Although our main focus is serine/threonine kinases (STKs)—where conformational heterogeneity and perturbation-driven population shifts are underexplored—we also applied our computational framework and nomenclature to ABL protein, a hallmark tyrosine kinase (TK) and drug target for chronic-myeloid-leukemia. Our computational workflow recaptured the canonical DFG_in_ ↔ DFG_out_ transition and, critically, resolved intermediate and multiple metastable substates within each DFG basin. This extends our ensemble nomenclature and sampling strategy from STKs to TKs, providing a fine-grained, multi-state view of kinase dynamics than the conventional two-state model.

The traditional research paradigm for kinases has primarily focused on retrospectively analyzing protein conformations^5^ and sampling conformational transitions between known end states. In contrast, the ability to prospectively sample the conformational heterogeneity in a diverse set of apo STKs with different gatekeeper residues and ability to capture population shifts and the kinetics associated with ensemble perturbations, allows for a deeper understanding of mechanistic insights. Our frameworks can be extended across a wide range of kinase subclasses and other dynamic protein families, such as G protein-coupled receptors (GPCRs)^23^ and tumor necrosis factors (TNFs)^24^, to gain mechanistic insights and identify druggable pockets. This framework can de-risk and accelerate early-stage drug discovery across a wide-range of targets.

The ability of our pipeline to sample “*rare*” or “*out-of-distribution*” conformational ensembles in biomolecules can be exploited as a data generation tool. Our study shows that generating high-quality data that captures a spectrum of conformational heterogeneity in biomolecules is key to gain generalizable mechanistic insights of molecular motion in protein classes.

## Results

### Defining Conformational Heterogeneity in Serine-Threonine Kinases Beyond DFG In/Out

Traditional classification of protein kinase structures has focused on clustering kinase macrostates into **DFG_in_** and **DFG_out_**conformations. This classification is based on the dihedral angle of the DFG-Phe sidechain rotamer and distance vectors involving the DFG motif and the αC-helix (**Figure 2A**). However, these nomenclature schemes overlook the conformational heterogeneity within the DFG_in_ ensemble. Using our state-of-the-art computational pipeline, we comprehensively sampled the full range of conformational heterogeneity within the DFG_in_ ensemble (detailed in the following sections). This enabled us to introduce a novel nomenclature scheme that fully captures the observed conformational diversity in serine-threonine kinases.

**Figure 2.**
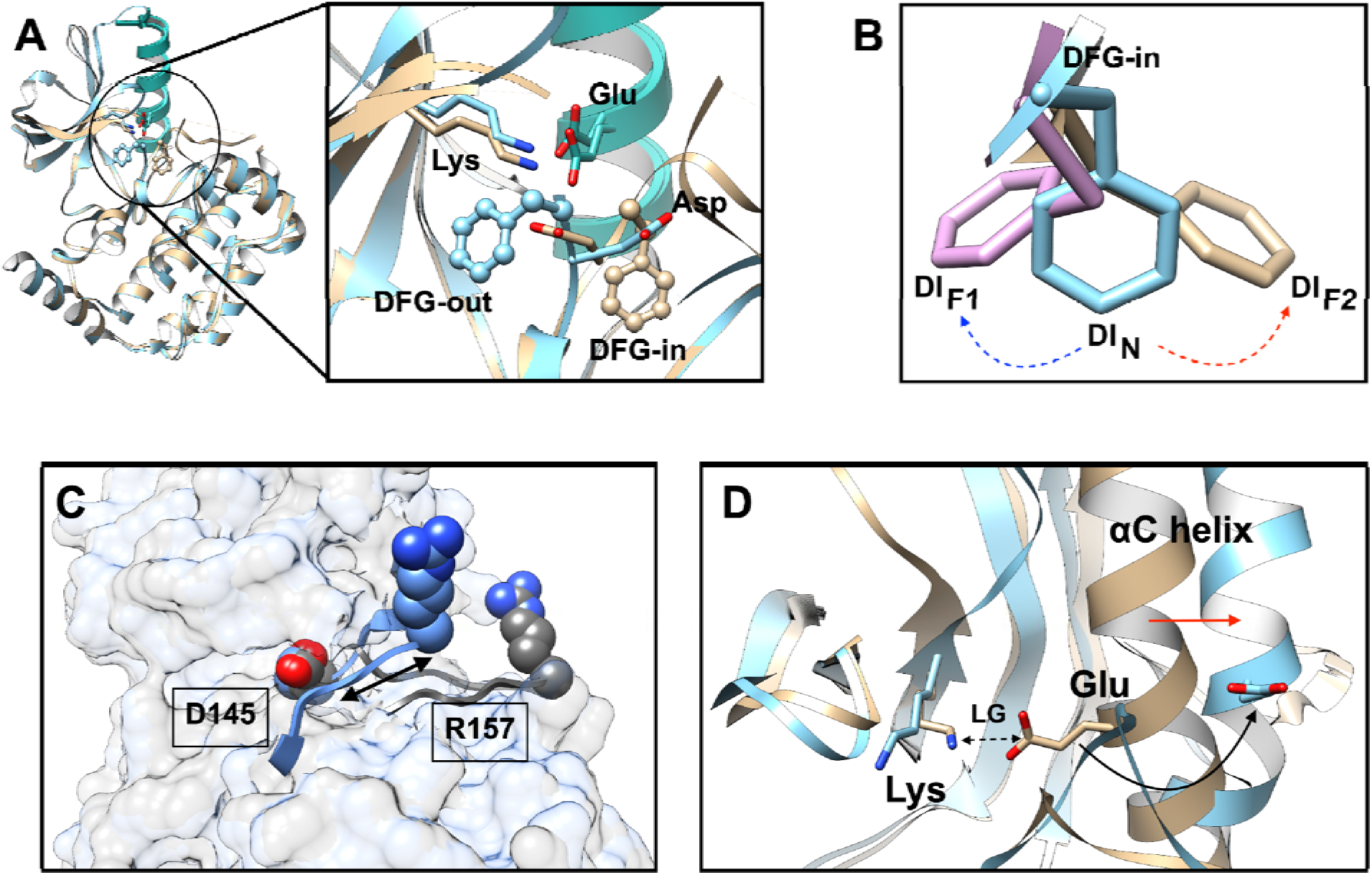
A: Pictorial representation of DFG_in/out_ snapshots in serine–threonine kinases, highlighting the positions of the catalytic DFG-Phe/Asp and conserved Lys and Glu residues. The *α*C-helix is shown in seagreen. **B:** Structural depiction of the DI_N/F1/F2_ conformations within the DFG_in_ macrostate. **C:** The activation loop of CDK2 is highlighted as ribbon, and the distance (*C*_*--C*_) between Asp145 and Arg157 (indicated by the arrow) is used to define the extended (AC_out_, grey) versus collapsed (AC_in_, blue) conformations. **D:** Definition of latched (LG_L_) versus unlatched (LG_U_) conformations involving Lys and Glu, underscoring the inward-to-outward shift of the *α*C-helix and the resulting changes in the relative positions of Lys and Glu.

### Ensemble Definition

We categorized the global DFG_in_ (**DI**) ensemble into three distinct states—**DI_N_**, **DI_F1_**, and **DI_F2_**—based on the χ angle (**Figure 2D**) of the conserved DFG-Phe side chain:

a. **DI_N_**: χ angle between –2 radians and 0 radians.
b. **DI_F1_**: χ angle between 0 radians and +2 radians.
c. **DI_F2_**: χ angle below –2 radians or above +2 radians.

### Lys-Glu Interaction in Serine-Threonine Kinases

Serine-threonine kinases are often characterized by a conserved salt-bridge interaction between a lysine residue in the β strand and a glutamate residue in the α-C helix (Figure 2A). To capture the heterogeneity of the Lys-Glu (**LG**) interaction, we introduce the following criteria:

1. **LG Latched (LG_L_)**: Distance between Lys-Nζ and Glu-δ < 0.45 nm.
2. **LG Unlatched (LG_U_)**: Distance between Lys-Nζ and Glu-δ > 0.45 nm.

The transition from **LG_L_** to **LG_U_**also reflects the α-C helix transitioning from an "in" to an "out" conformation (**Figure 2B**). The macrostates defined by **DI_N_**, **DI_F1_**, and **DI_F2_** can each encompass multiple microstates, where the Lys-Glu interaction remains either latched or unlatched. These microstates are: **DI_N_LG_L_, DI_N_LG_U_, DI_F1_LG_L_, DI_F1_LG_U_, DI_F2_LG_L_, DI_F2_LG_U_** .

### Activation loop dynamics in Serine-Threonine Kinases

The activation loop in serine/threonine kinases (STKs) plays a crucial role in molecular recognition and exists in a dynamic equilibrium between extended and collapsed conformations^25^. We capture the conformational heterogeneity of the activation loop by measuring the distance between a stable anchor point (DFG-Asp) and the conserved arginine residue within the activation loop (**Figure 2C**). The specific residue numbers for the anchor point and arginine are highlighted in the Supporting Information. This measurement allows us to define the conformational states as **AC_out_** (extended) and **AC_in_** (collapsed). When combined with the conformational states defined by **DI_N/F1/F2_** and **LG_U/L_**our approach effectively captures the full spectrum of conformational heterogeneity of the activation loop in STKs.

### Incorporating DFG_out_ states

Our ensemble definitions can be extended to include DFG_out_ (**DO**) and other intermediate ensembles (characterized by the distance between Glu-CA—Phe-CZ and Lys-CA—Phe-CZ), while preserving all other microstate definitions associated with χ angle of DFG-Phe and Lys– Glu (**LG**) interactions and the dynamics of activation loop (**AC**).

Since STKs remain in the **DI** ensemble in the absence of a type-II inhibitor, we will focus our discussion on the conformational heterogeneity of the **DI** ensemble and how perturbations influence the thermodynamics and kinetics of various metastable states. Additionally, we will highlight how our ensemble definition can be extended to incorporate transition between DFG_in_ and DFG_out_ states as well novel intermediate states in a tyrosine kinase which captures full range of conformational heterogeneity.

### V600E mutation and Ligand Binding Modulate BRAF Conformational Dynamics and BRAF–MEK1 Interactions

The interaction between the serine-threonine kinase BRAF and MEK1 is a key step in the RAS– RAF–MEK–ERK signaling cascade, which regulates essential cellular processes such as growth, proliferation, and differentiation. BRAF is mutated in 66% of malignant melanomas, with the single-point mutation V600E accounting for approximately 80% of these cases. This BRAF V600E mutation results in a permanently active BRAF protein, continuously activating the MAPK pathway and driving cancer cell proliferation^26–28^. However, the specific effects of V600E on the conformational dynamics of both the BRAF–MEK1 complex and the BRAF monomer remain an open question.

Physics-based molecular simulations of the wild-type BRAF–MEK1 complex, starting from an AlphaFold3-predicted structural ensemble, reveal that the complex predominantly adopts an inactive conformation, existing in a dynamic equilibrium between the **DI_F1_** (78.71%) and **DI_F2_**(21.29%) states (**Figure 3**). This inactive state also features Lys-Glu in unlatched conformations (**DI_F1_LG_U_**, **DI_F2_LG_U_**). In contrast, the V600E mutation shifts the population toward the active state (**Figure 4**, **Figure S2** in Supporting Information), evidenced by an increased population of the **DI_N_**state (75.91%) compared to **DI_F1_** (15.42%) and **DI_F2_**(8.67%). Moreover, molecular dynamics simulations capture conformational heterogeneity within the **DI_N_** state in V600E BRAF–MEK1, which transitions between **DI_N_LG_L_**and **DI_N_LG_U_** conformations (**Figure S1** in Supporting Information).

**Figure 3.**
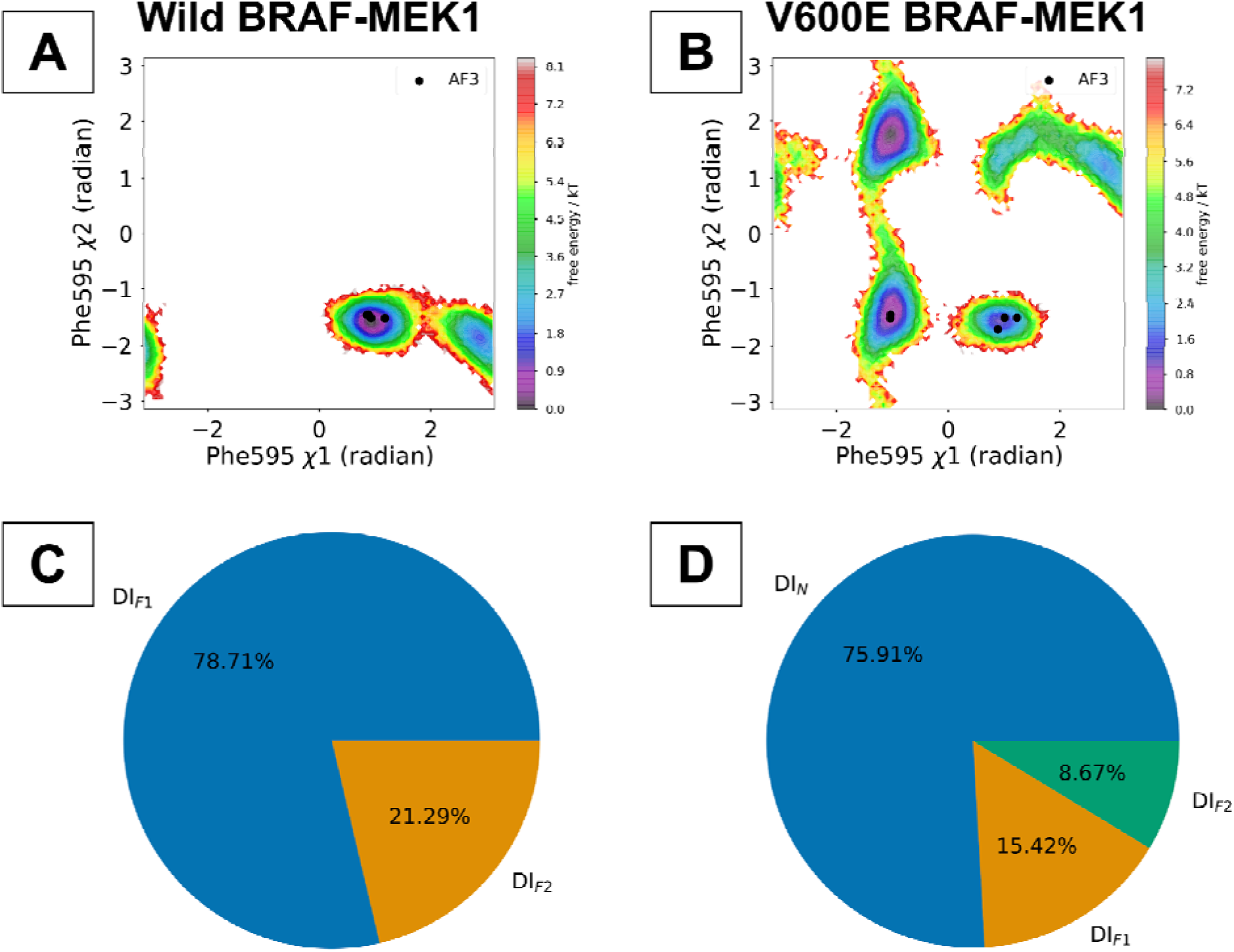
Markov state modeling (MSM) based on AI-augmented molecular simulations, initiated from an AlphaFold3-derived conformational ensemble, reveals conformational heterogeneity in the BRAF– MEK1 complex. In the wild-type complex, BRAF remains in a dynamic equilibrium between DI_F1_ and DI_F2_ states (A, C). However, the V600E mutation shifts the population from DI_F1/F2_ to the DI_N_ state (B, D). Although AlphaFold3 can sample single structures corresponding to DI_N_ and DI_F1/F2_ conformations in V600E BRAF–MEK1, it does not capture the population shift upon mutation, emphasizing th importance of physics-based molecular simulations for a comprehensive characterization of conformational heterogeneity.

**Figure 4.**
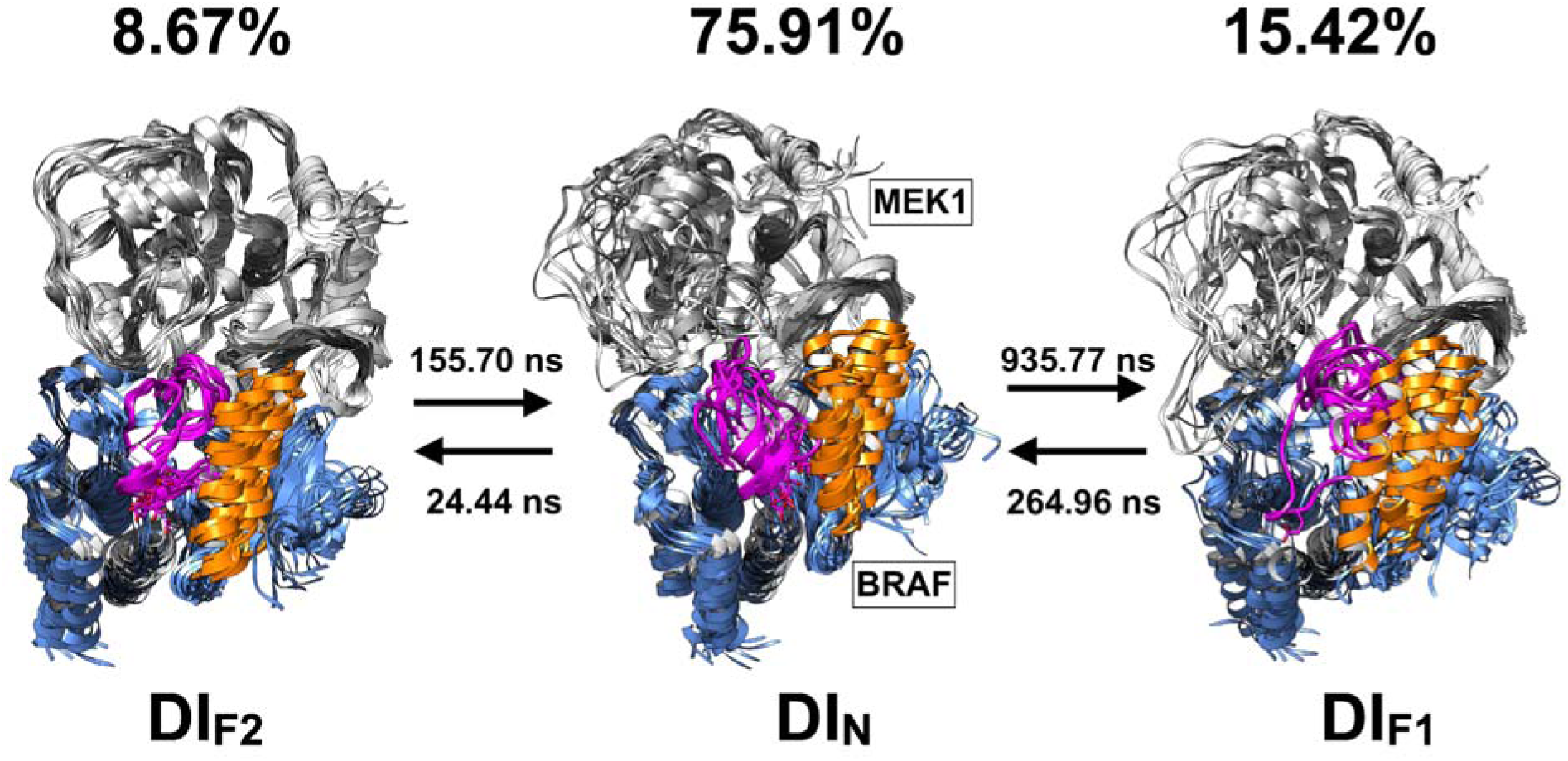
Timescales associated with conformational transitions in V600E BRAF–MEK1 complex reveal a faster transition from DI_F1_ to DI_N_ (264.96 ns) than from DI_N_ to DI_F1_ (935.77 ns). In contrast, wild-type BRAF–MEK1 remains in a dynamic equilibrium between DI_F1_ and DI_F2_. Furthermore, the V600E BRAF– MEK1 complex exhibits conformational heterogeneity in the *α*C-helix (orange) and the activation loop (magenta), transitioning from an inward (DI_N_) to outward (DI_F1_) orientation in the *α*C-helix and from an extended (DI_N_) to a collapsed (DI_F1_) state in the activation loop.

Predicting the structural ensemble of the **DI_F1_** conformation in the V600E BRAF–MEK1 complex enabled the identification of a hydrophobic backpocket in MEK1. Targeting this pocket with small molecules broadens the scope for designing novel therapeutic modalities that could stabilize the BRAF-MEK1 complex in the **DI_F1_**conformation, opening a promising avenue for developing selective treatments against BRAF V600E–mutated melanoma (**Figure 5**).

**Figure 5.**
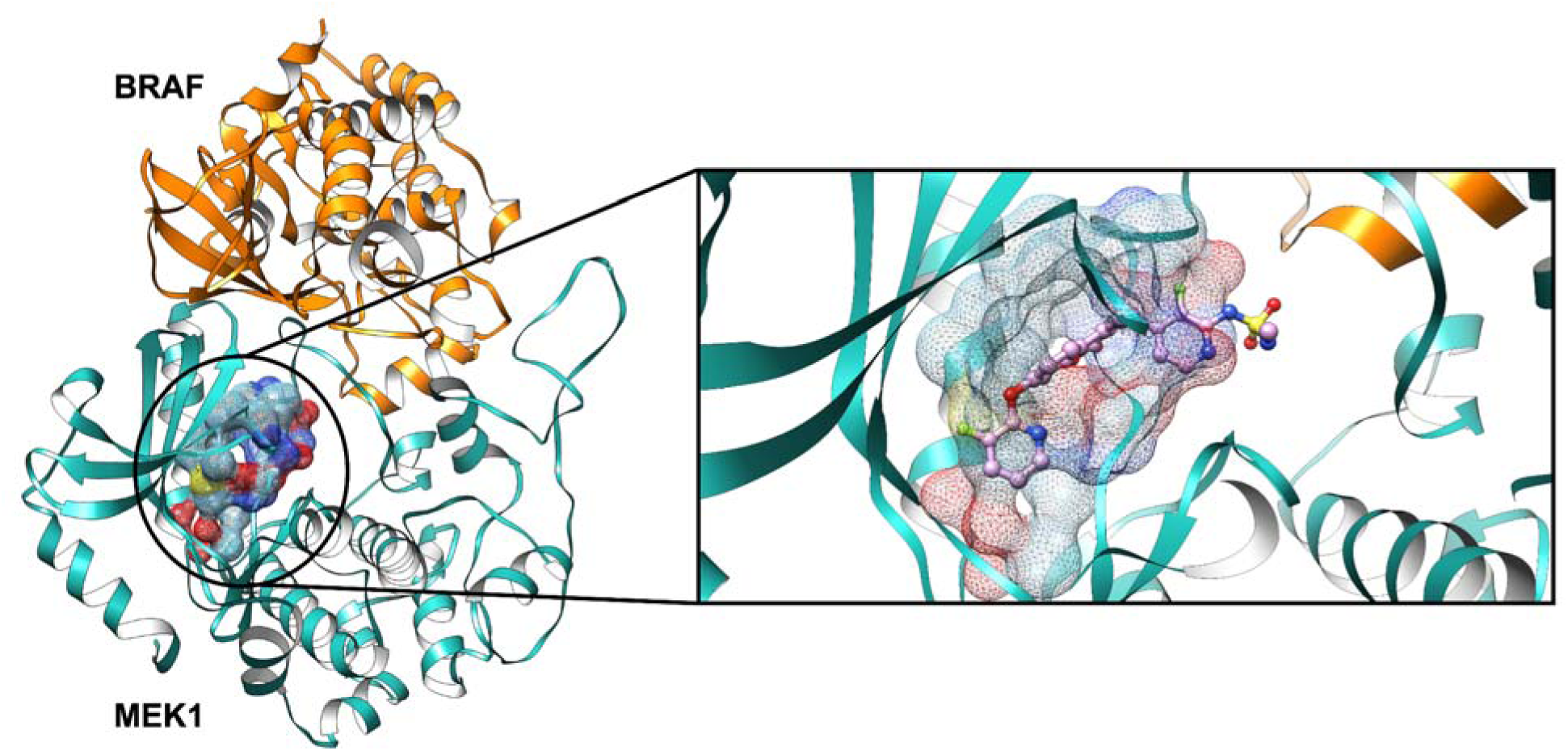
A structural snapshot from the conformational ensemble of the V600E BRAF–MEK1 complex reveals a hydrophobic backpocket in MEK1 with a druggability score of 0.880 predicted by FPocket^29^. Targeting this pocket with novel therapeutics can stabilize the V600E BRAF–MEK1 complex in the **DI_F1_**state, potentially restoring normal cellular function. Notably, the panRAF–MEK1 molecular glue NST628 (PDB: 9AXX)^30^, shown in ball-and-stick representation, binds to this hydrophobic backpocket as captured by our simulation.

Similar to the BRAF–MEK1 complex, the BRAF monomer also exhibits a population shift from the inactive (**DI_F1/F2_**) to the active (**DI_N_**) state upon V600E mutation (**Figure S4** in Supporting Information). Vemurafenib^31^ was discovered as a highly specific, ATP-competitive inhibitor of BRAF V600E with selectivity against melanoma cells; however, how vemurafenib affects the conformational dynamics of V600E BRAF remains an open question. Molecular dynamics simulations initiated from V600E BRAF bound to vemurafenib in the **DI_N_** state capture a shift in population towards the inactive state (**Figure 6** and **Figure S5** in Supporting Information), reflected by an increased population of the **DI_F1_**and **DI_F2_** states in the vemurafenib–V600E BRAF complex (**DI_F1_**: 23.50%, **DI_F2_**: 2.40%) compared to the apo V600E BRAF (**DI_F1_**: 12.86%, **DI_F2_**: 0.02% ).

**Figure 6.**
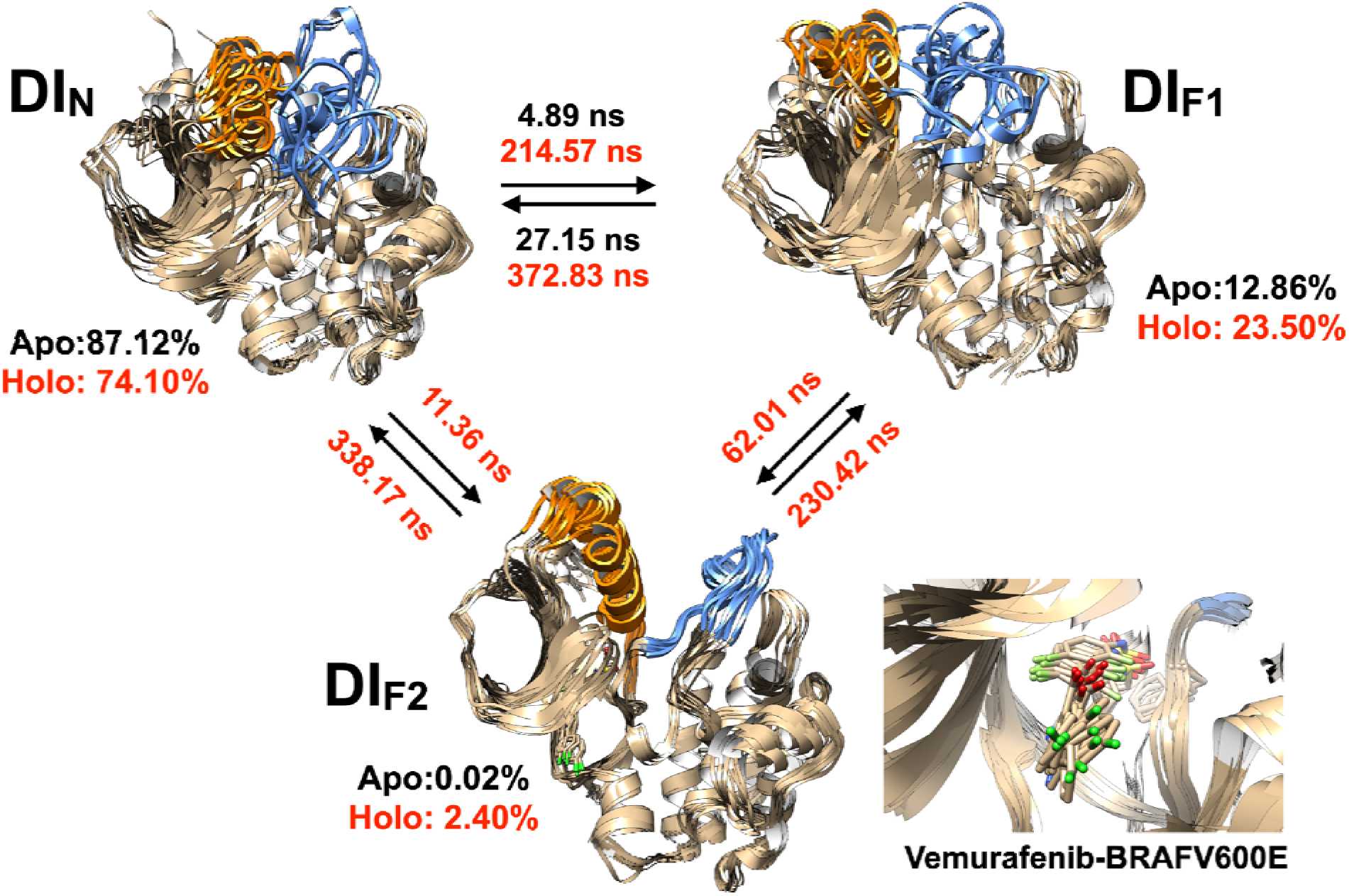
AI-augmented molecular simulations and ML-augmented adaptive sampling comprehensively capture the full range of conformational dynamics in the apo V600E BRAF monomer and th vemurafenib-bound holo V600E BRAF monomer respectively. Vemurafenib binding shifts the ensembl from DI_N_ to DI_F1/F2_, as evidenced by faster DI_N_ → DI_F1_ (214.57 ns) and DI_N_ → DI_F2_ (11.36 ns) transitions compared to DI_F1_ → DI_N_ (372.83 ns) and DI_F2_ → DI_N_ (338.17 ns) in the holo form. Notably, the DI_F2_ state in apo V600E BRAF is sampled too sparsely to predict transition times via PCCA+^32^. The αC-helix and activation loop display marked conformational heterogeneity, highlighted in orange and blue, respectively.

Conformational transitions from the **DI_N_**state to the **DI_F1_** state increase the population of **DI_F1_LG_U_** conformation in holo state (16.93%) compared to apo V600E BRAF (7.96%), a hallmark of active-to-inactive transitions in serine-threonine kinases (**Figure S1** in Supporting Information).

### Conformational Heterogeneity and Allostery in RIPK2: How Mutations and Protein-Protein Interactions Induce Population Shifts

The interaction between the kinase domain of receptor-interacting protein kinase 2 (RIPK2)—a serine-threonine kinase—and the E3 ligase X-linked Inhibitor of Apoptosis Protein (XIAP) serves as a crucial checkpoint in regulating inflammation and innate immunity, making RIPK2 a potential therapeutic target for diseases characterized by excessive or chronic inflammation^33–35^. A K47R mutation in RIPK2 alters its conformational state^36^. However, relying on a single static structure cannot adequately capture the dynamic nature of RIPK2, nor how mutations and protein–protein interactions modulate its conformational landscape.

To address this gap, a Markov state model (MSM) trained on AlphaFold-seeded molecular simulations was used to identify the populations and timescales associated with RIPK2’s conformational states. In the wild-type protein, RIPK2 predominantly occupies the **DI_N_** state (88.63%), with smaller populations in **DI_F1_** (10.45%) and **DI_F2_** (0.92%). Previously, **DI_N_** wa defined as the active state, while **DI_F1_**and **DI_F2_**were categorized as inactive states. In the K47R mutant, the conformational equilibrium is shifted toward the inactive states, with the population of **DI_F1_** and **DI_F2_** predicted to be 24.00% and 2.82% respectively (**Figure 7** and **Figure S7** in Supporting Information).

**Figure 7.**
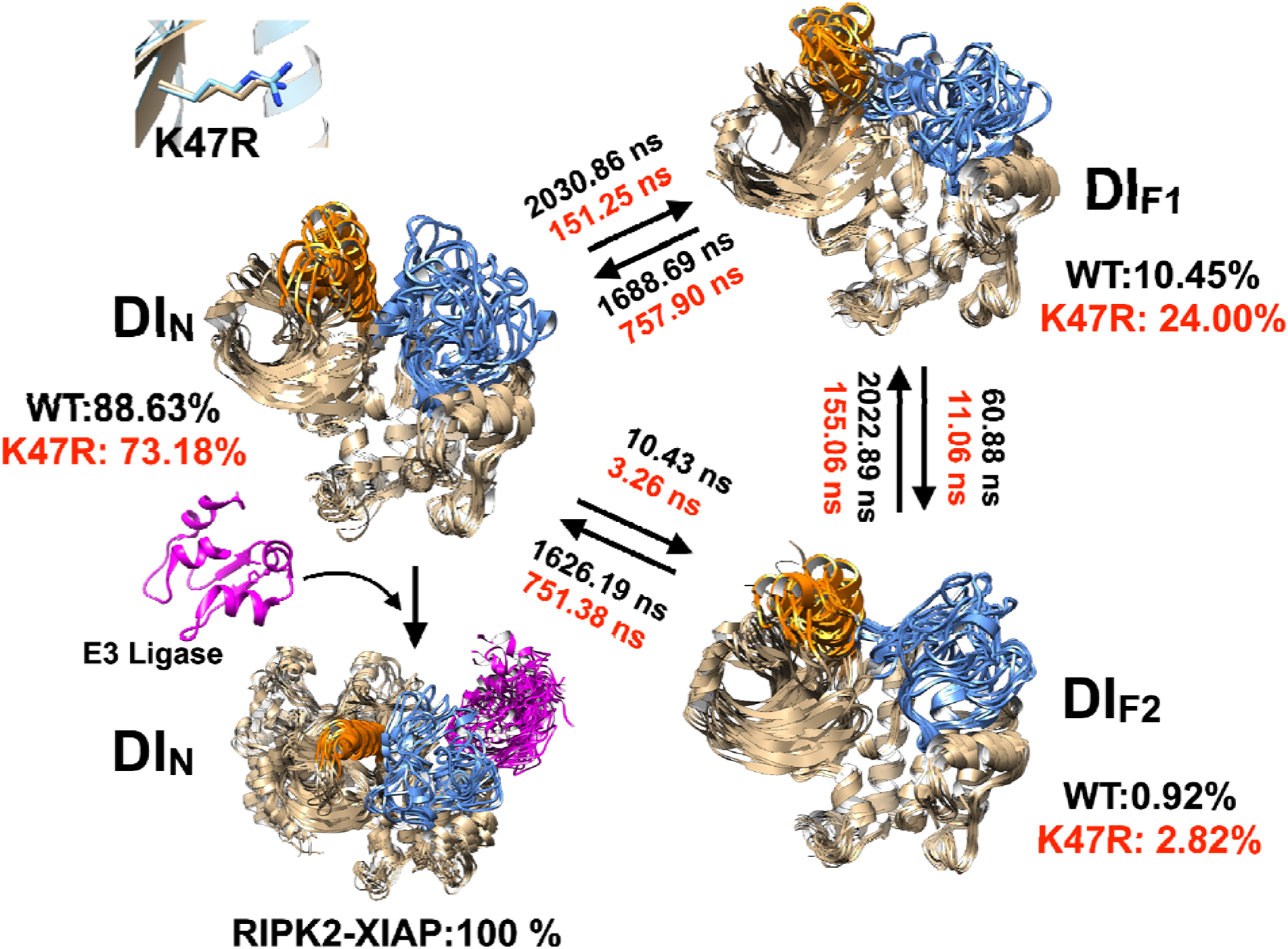
Populations and timescales associated with conformational transitions indicate that RIPK2 remains in a dynamic equilibrium among the **DI_N_**, **DI_F1_**, and **DI_F2_**states. In wild type apo RIPK2, th active (**DI_N_**) state predominates. However, the K47R mutation shifts the equilibrium toward the inactiv (**DI_F1_** and **DI_F2_**) states. This population shift is reflected in the mean first-passage transition times, showing that the K47R mutation accelerates the **DI_N_** → **DI_F1/F2_** transitions compared with wild-type RIPK2. By contrast, binding of the E3 ligase XIAP stabilizes the **DI_N_** state. The ability to predict conformational heterogeneity and understand how perturbations—such as mutations and XIAP binding— modulate RIPK2 dynamics is critical for capturing molecular recognition and developing selective therapeutics targeting *active-to-inactive* transitions. Conformational ensembles corresponding to each state capture the heterogeneity of the *α*C-helix (orange) and the activation loop (blue).

Additional physics-based molecular simulations also captured the conformational heterogeneity of microstates. In wild-type RIPK2, most of the population in the **DI_N_** state remains in the Lys-Glu latched conformation (**LG_L_**) rather than the unlatched (**LG_U_**) conformation. Transitioning from **DI_N_** to **DI_F1_** shifts the population towards the unlatched conformation. This conformational change from **DI_N_LG_L_** to **DI_F1_LG_U_**involves a shift of the *αC-helix* from an “in” to an “out” conformation—a key hallmark of kinase conformational transitions (**Figure S8** in Supporting Information). Transition of *αC-helix* from in to out opens up the allosteric backpocket in RIPK2 which can be targeted by type III inhibitors^37^ (**Figure 8**). It is key to highlight that AlphaFold alone failed to sample structures corresponding to **DI_F1/F2_**conformations in wild type RIPK2.

**Figure 8.**
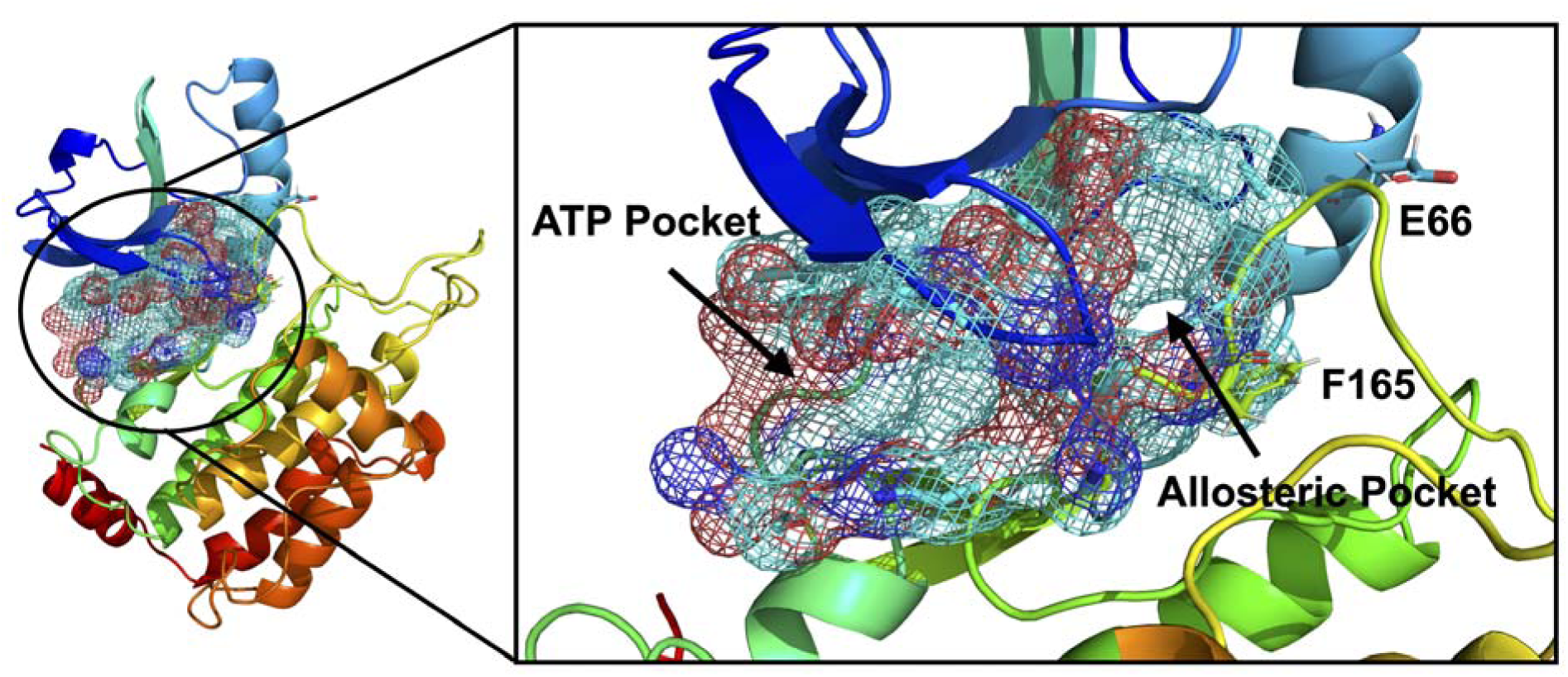
Pictorial representation of the DI_F1_LG_U_ conformational snapshot of apo RIPK2, illustrating the allosteric backpocket that can be targeted by type III kinase inhibitors. The formation of this backpocket is driven by the DI_N_ -to-DI_F1_ flip and the transition of the α*C-helix* from the “ in” to “out” state.

Our time-lagged autoencoder based machine learning model (ML model) trained on high-dimensional temporal data from molecular dynamics simulations of RIPK2 identifies slowly varying structural features governing its conformational dynamics. The latent layers of the ML model serve as a two-dimensional collective variable—specifically, a nonlinear combination of dihedral angles—used to construct MSM (**Figure S9** and **S10** in Supporting Information). The model detects the flipping of Trp170 and Ile208. Flipping of Trp170 signifies a transition of the activation loop from an active to an inactive conformation as highlighted by *Vats et al*^21^, whereas flipping of Ile208 alters the RIPK2–XIAP binding interface and disrupts the hydrophilic interaction involving Lys209 in RIPK2 and Asn209 and Glu211 in XIAP (**Figure 9**).

**Figure 9.**
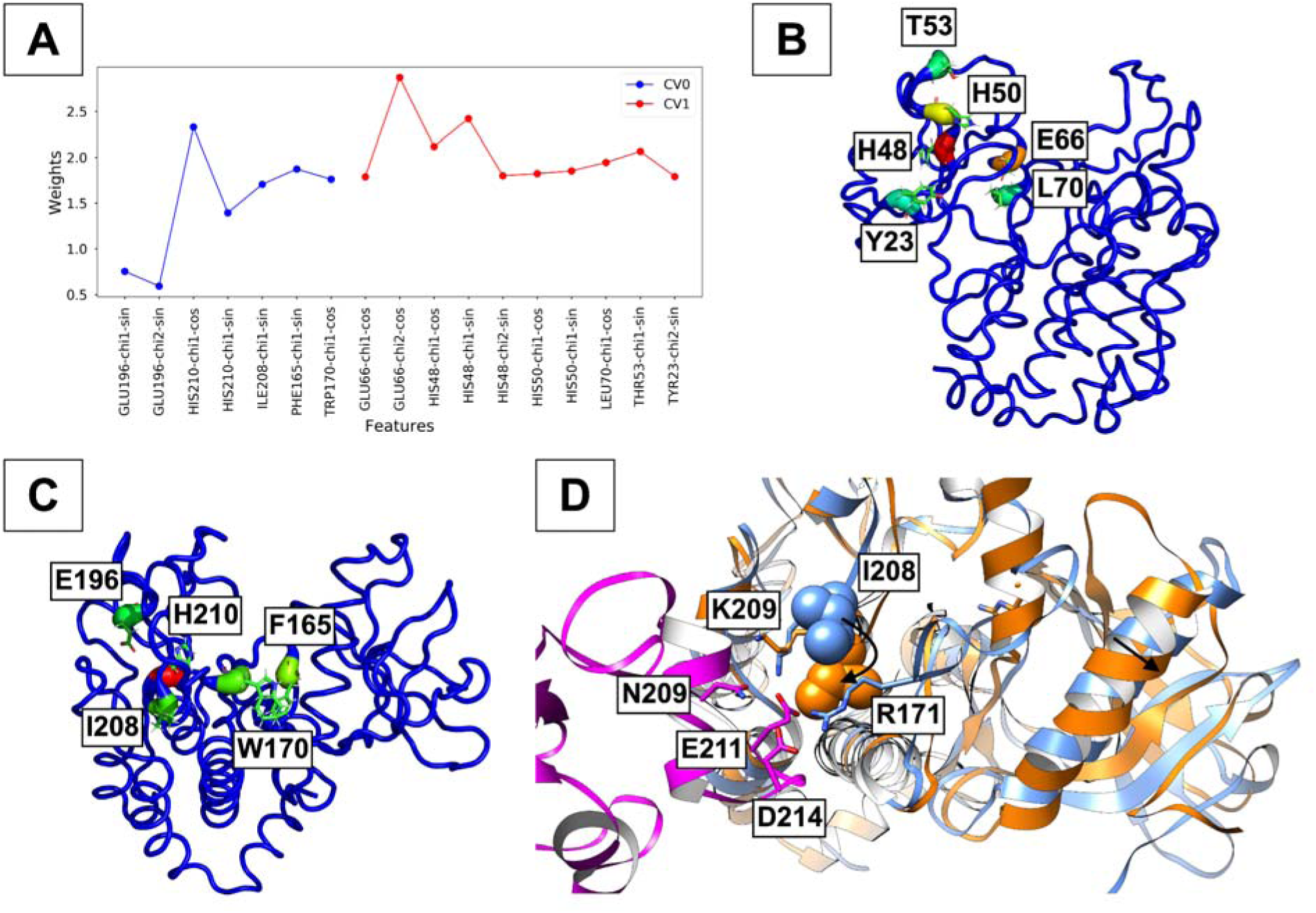
Latent layers of the ML model capture key structural features of RIPK2 (A). Specifically, CV0 reflects the C-lobe dynamics (defined by the χ1 and χ2 angles of residues 164–215), while CV1 encompasses the N-lobe dynamics (defined by the χ1 and χ2 angles of residues 14–74). ML weights projected onto the structural features reveal important conformational motions governing both N-lobe (B) and C-lobe (C) dynamics of RIPK2. In particular, the model highlights critical residues governing conformational heterogeneity of the αC-helix and the activation loop, which underpin transitions between RIPK2’s active and inactive states. Within the activation loop, flipping of Trp170 disrupts the hydrogen bond between Arg171 of RIPK2 and Asp214 of XIAP. Simultaneously, flipping of Ile208 interrupts the interaction between Lys209 of RIPK2 and Asn209/Glu211 of XIAP (D). Together, these events illustrat the molecular basis of conformational allostery, showing how RIPK2 transitions from the active (blue) to the inactive (orange) state and how these changes modulate RIPK2–XIAP interactions.

Furthermore, MSM-predicted transition times indicate that the K47R mutation accelerates conformational transitions between the **DI_N_** ↔ **DI_F1_**and **DI_N_**↔ **DI_F2_** states compared to wild-type RIPK2. Specifically, the mean first-passage time (MFPT) for the **DI_N_**→ **DI_F1_** transition i ∼2030 ns in the wild type and ∼151 ns in the mutant, while the **DI_N_** → **DI_F2_** transition is ∼10 ns in the wild type and ∼3 ns in the mutant. A similar trend is observed for the reverse **DI_F1/F2_** → **DI_N_**transitions, highlighting that the K47R mutation shifts the population and the timescales of these conformational transitions towards **DI_F1/F2_**states which shifts the dynamics of the activation loop from extended (**AC_out_**) to collapsed (**AC_in_**) conformations (**Figure 7 and Figure S11 and S12 in Supporting Information**).

Unlike K47R mutation, RIPK2 and XIAP interaction stabilizes the **DI_N_** state (**Figure 7** and **Figure S7** in Supporting Information). Taken together, these results demonstrate how both mutation and protein–protein interactions modulate the conformational dynamics of RIPK2, extending beyond the static structure paradigm of X-ray crystallography and AI-based predictions such as AlphaFold.

### Conformational Dynamics of Serine-Threonine Kinases: A Generalizable Model

Markov state models (MSMs) enabled us to capture the populations and transition times of **DI_N_**, **DI_F1_**, and **DI_F2_**states in an expanded set of serine-threonine kinases, including MAPK14^38^, IRAK1 (Interleukin-1 receptor-associated kinase 1), IRAK4 (Interleukin-1 receptor-associated kinase 4)^39^, and CDK2 (Cyclin-Dependent Kinase 2) ^25^. In all of these proteins, the apo protein maintains a dynamic equilibrium among **DI_N_**, and **DI_F1/F2_**conformations (**Figure 10**). Population-weighted projections along DFG-Phe χ1 and Lys–Glu distances further revealed **LG_L/U_** microstate heterogeneity within these **DI_N/F1/F2_** macrostates (**Figure 11**).

**Figure 10.**
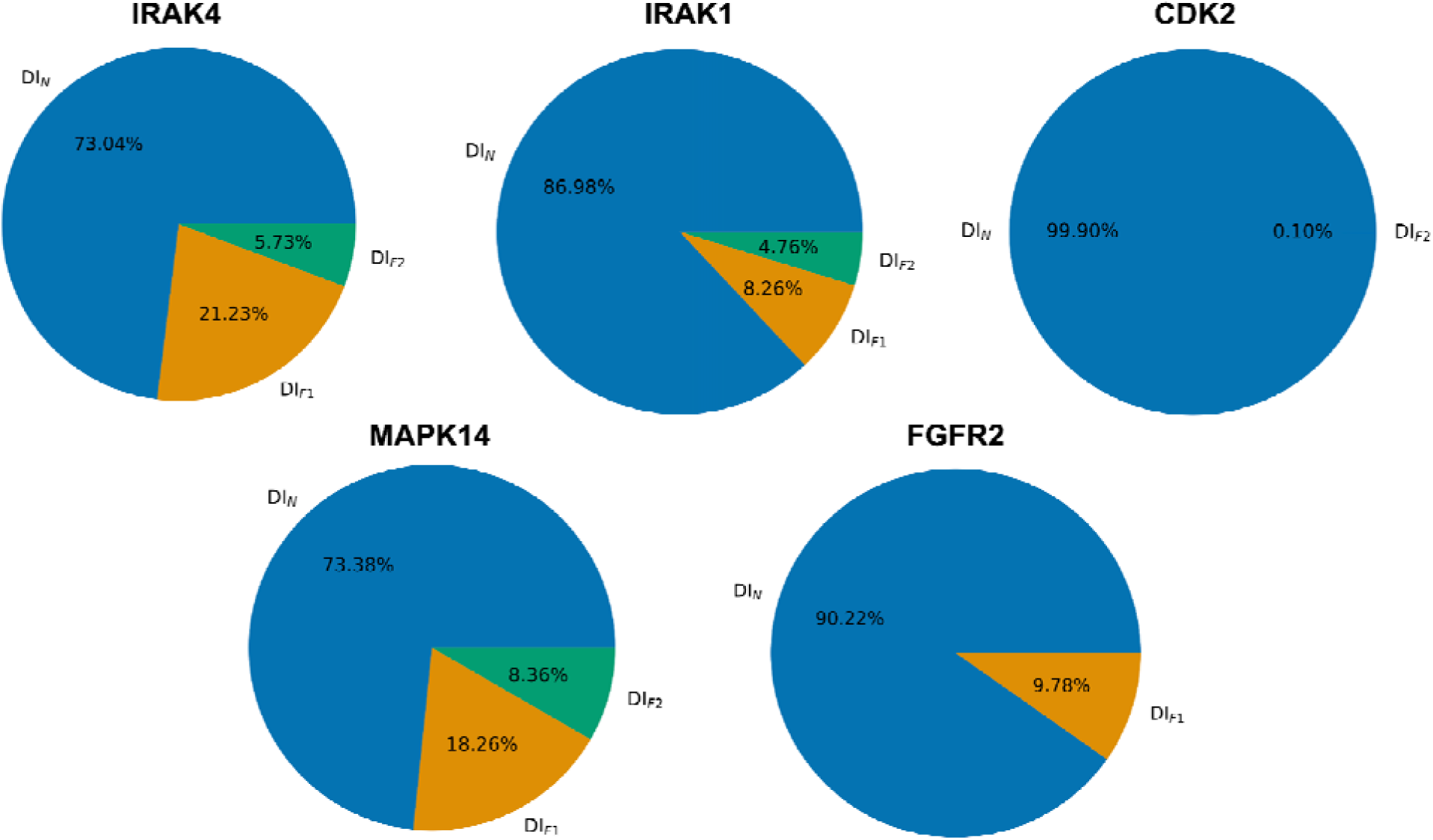
MSMs trained on physics-based molecular simulations of multiple serine-threonine kinases (IRAK1/4, CDK2, and MAPK14) and the tyrosine kinase FGFR2^40^ reveals generalizable conformational dynamics, showing that kinases remain in dynamic equilibrium between DI_N_ and DI_F1/F2_ states with variable populations.

**Figure 11.**
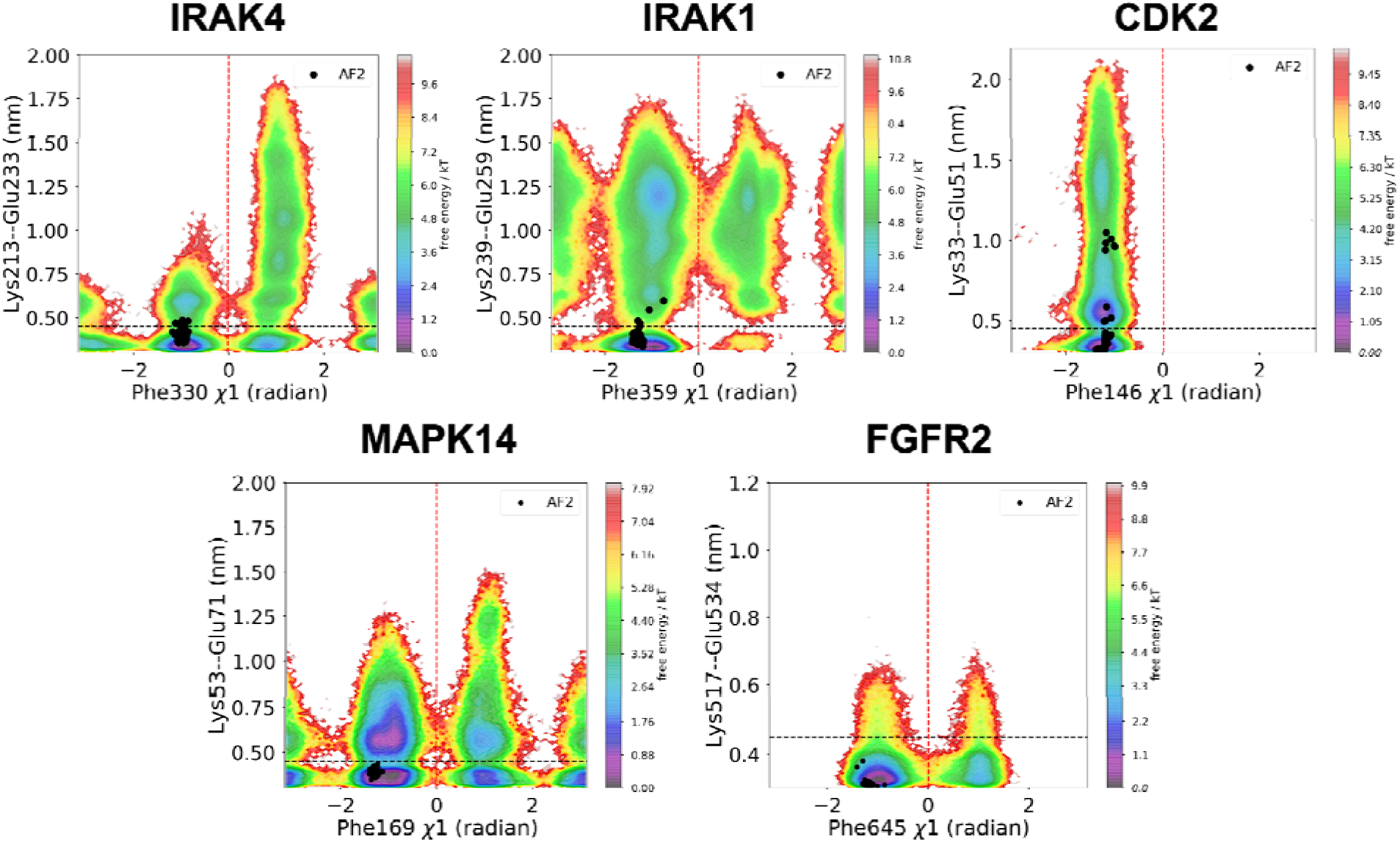
The population-weighted free energy surface derived from the MSM, projected along the χ1 angle of DFG-Phe and the Lys–Glu distance, reveals conformational heterogeneity defined by **LG_L/U_**microstates within the broader **DI_N/F1/F2_** macrostates. However, in all cases, conformational ensemble generated by MSA subsampling fail to capture the full spectrum of this heterogeneity.

Notably, in IRAK1, AlphaFold-seeded molecular dynamics simulations only partially sampled the conformational space defined by the **DI_F1_**, and **DI_F2_** states. However, adaptive sampling initiated from conformational snapshots obtained through our ML model’s latent layer captured the full spectrum of conformational heterogeneity (**Figure S13** in Supporting Information).

Conformational variability in kinases extends beyond the DI_N/F1/F2_ macrostates. For example, in the apo form of CDK2, the DI_N_ macrostate predominates but coexists in a dynamic equilibrium between AC_out_ (extended activation loop) and AC_in_ (collapsed activation loop) states involving the activation loop. Our ensemble definitions also capture conformational heterogeneity in the Lys-Glu interaction across AC_out/in_ ensembles (AC_out/in_LG_L/U_). Cyclin E1 binding stabilizes the ACout conformation (**Figure 12** and **Figure S14** in the Supporting Information). Although Cyclin E1 has long been considered undruggable, a high-resolution dynamic representation of the CDK2–Cyclin E1 complex—encompassing its full conformational heterogeneity—reveal cryptic binding pockets within the protein–protein interface, opening new avenues for targeting Cyclin E1 (**Figure 13**).

**Figure 12.**
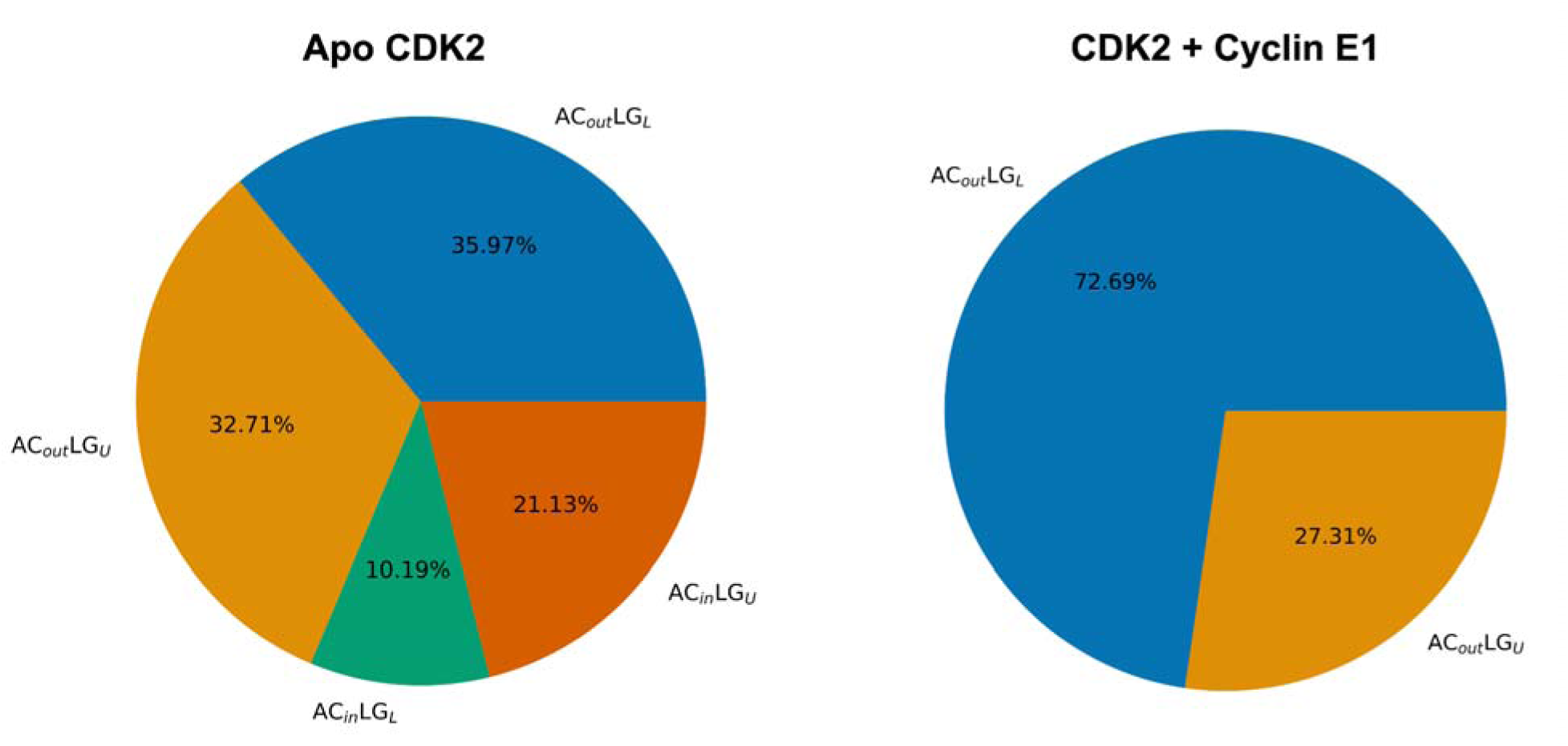
Our ensemble definition captures conformational heterogeneity of the activation loop and Lys-Glu interactions in apo CDK2 and CDK2-Cyclin E1 complex. Apo CDK2 remains in a dynamic equilibrium between AC_out_ and AC_in_ conformations which captures differentiable dynamics of the Lys-Glu interactions. Binding of Cyclin E1 stabilizes AC_out_ confirmation where the AC_out_ confirmation captures the dynamic equilibrium between LG_U_ and LG_L_ conformations.

**Figure 13.**
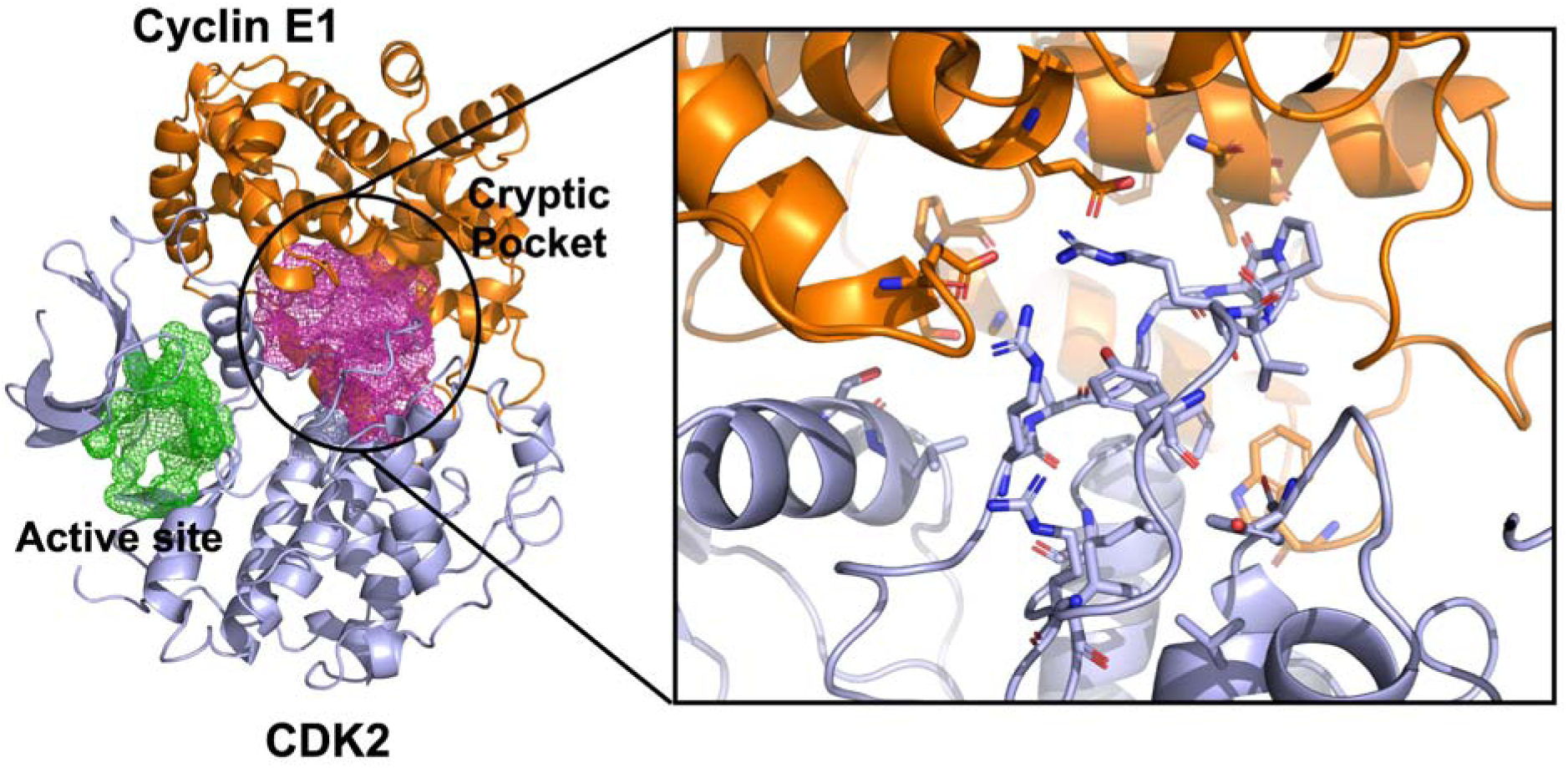
A structural snapshot from the DI_N_AC_out_ conformational ensemble of the CDK2–Cyclin E1 complex reveals a cryptic pocket at the protein–protein interface with a high druggability score (0.967) predicted by FPocket. Targeting this cryptic pocket provides an alternative approach to modulating the CDK2–Cyclin E1 complex beyond the traditional CDK2 active site.

Finally, the conformational heterogeneity captured by our computational framework is not exclusive to serine-threonine kinases. Our adaptive sampling framework also enabled DI_N_ to DI_F1_ conformational transitions in receptor tyrosine kinase Fibroblast Growth Factor Receptor 2 (FGFR2) demonstrating the generalizability of our ensemble-definition method for sampling conformational dynamics across the human kinome (**Figure 14**). It is notable that ultra-long simulations initiated from the apo FGFR2 structure (PDB: 1GJO) on specialized Anton hardware, as well as conformational ensembles generated through MSA subsampling and AF seeded MD simulations (Figure S14 in Supporting Information), fail to capture DI_N_ → DI_F1_ conformational transitions (**Figure 14 C, D**). In contrast, binding of the highly selective small molecule RLY-4008^40^ to FGFR2 stabilizes the DI_F1_ conformational state, underscoring the importance of prospectively capturing kinase conformational transitions for the development of selective therapeutics.

**Figure 14.**
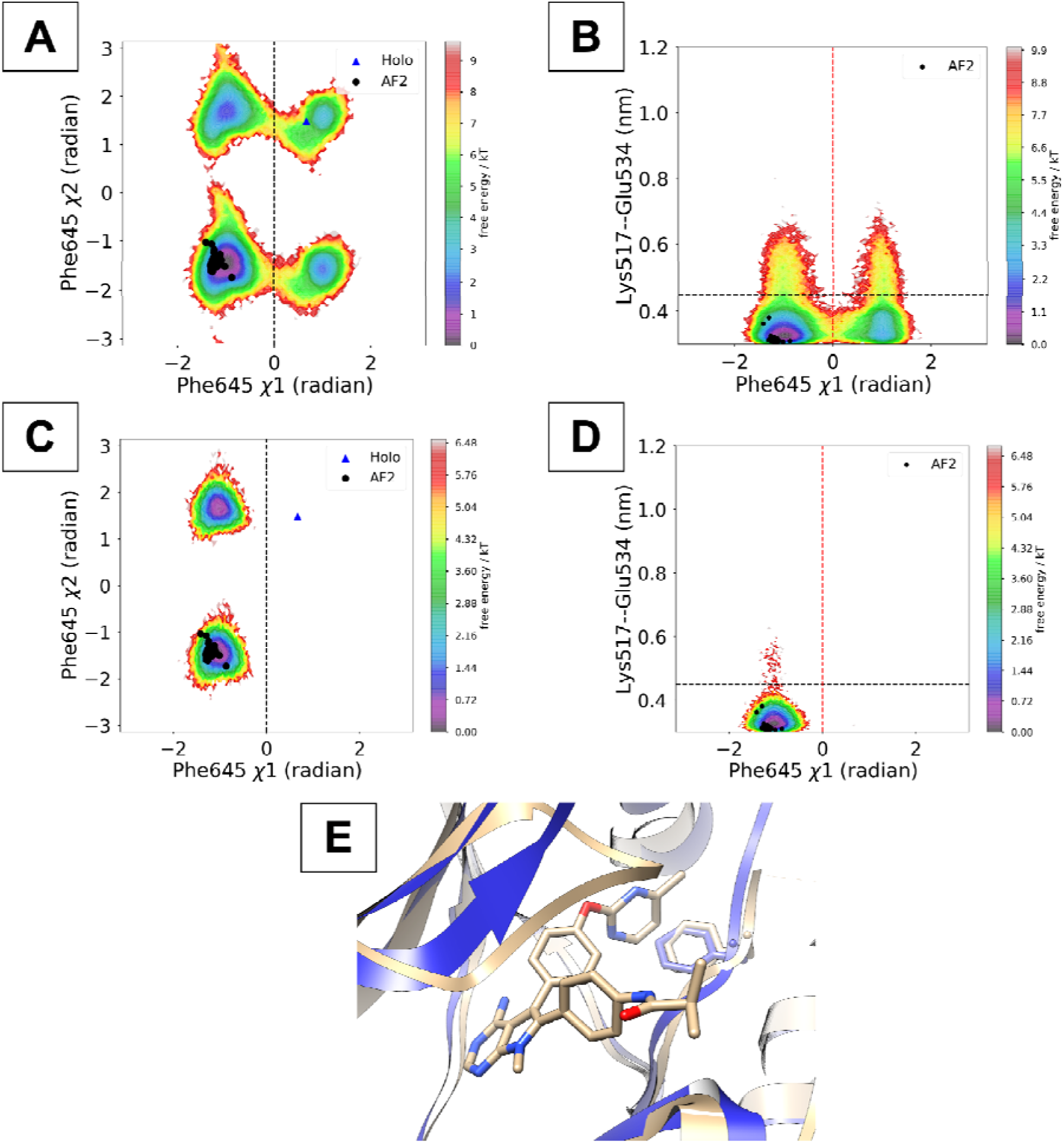
A population-weighted free energy surface, projected along DFG-Phe χ1/χ2 angles and th distance between DFG-Phe χ1 and Lys-Glu, illustrates how molecular simulation (100 replicas × 100 ns each, totaling 10 μs) initiated from physics-refined conformational ensemble captures apo FGFR2 conformational transitions (A, B) compared to ultralong MD simulations (three 25 μs replicas) performed on Anton. The blue triangle marks the holo FGFR2 structure (RLY-4008 + FGFR2 complex, PDB: 8STG^40^), while black dots represent conformational ensembles generated via MSA subsampling. A structural superimposition of the DI_F1_ state (blue) on holo FGFR2 (PDB: 8STG, silver) highlights the DFG-Phe conformation and the relative positioning of the small molecule (E).

### AI accelerated molecular simulation captures DFG_in_ to DFG_out_ transitions in apo tyrosine kinase

Our physics-based molecular simulations, initiated from an AlphaFold-generated conformational ensemble of apo wild-type ABL kinase, successfully captured the transition between DFG_in_ and DFG_out_ states in a total of 8 microseconds of trajectory. This represents a dramatic acceleration compared to previously published work, which required 10 *milliseconds* of trajectory to observe the same transition. The majority of the population in DFG_in_ state remains in Phe_N_ (analogous to DI_N_ state) state defined by DFG-Phe χ angle between –2 radians and 0 radians. Transition from DFG_in_ to DFG_out_ shifts the population towards Phe_F2_ state (analogous to DI_F2_ state) defined by DFG-Phe χ angle below –2 radians or above +2 radians. This populations shift can be also reflected by increased population of LG_U_ conformations upon DFG_in_ to DFG_out_ transitions in apo ABL kinase. This is consistent with previous observations where transition to DFG_out_ state pushes the αC-helix to ‘*out*’ conformation which opens up an extended backpocket that can be targeted by type-II kinase inhibitors.

Recent NMR experiments have illuminated the conformational transitions and intermediate states within the DFG_in_ and DFG_out_ ensembles. Our approach not only captured these known transitions but also sampled two novel conformational ensembles: DFG_Neo1_ and DFG_Neo2_. DFG_Neo1_ is characterized by a hydrogen bond interaction between Lys290 and DFG-Asp, which pulls the *P-loop* closer to the DFG residues, while Phe401 (DFG-Phe) adopts a DFG_out_ state. DFG_Neo2_ captures transient conformational ensemble, capturing the transition states between DFG_in_ and DFG_out_. We believe stabilizing DFG_Neo1_ via small molecules offers a path to developing next-generation therapeutics for imatinib-resistant leukemia.

A key challenge in kinases is to capture long-timescale conformational dynamics of apo proteins which governs functionally important conformational transitions. Time-lagged autoencoder trained on dihedral angles derived from AlphaFold seeded molecular simulation of ABL kinase, provides a sophisticated latent representation of the conformational dynamics, allowing us to accurately estimate unbiased population shifts and autonomously pinpoint the key residues (**Figure S19** in Supporting Information) that control these dynamic processes.

## Discussion

Serine/threonine kinases exist in a dynamic equilibrium among multiple metastable states. The conformational heterogeneity of these states governs their function and is critical for developing selective therapeutics. We introduced ensemble definitions to capture the full spectrum of conformational heterogeneity and employed AI-augmented molecular simulations and adaptive sampling to examine how the dynamical exchange among these ensembles is modulated by protein–protein interactions, small-molecule binding, and disease-causing mutations.

While the classic view of protein kinases has centered on the transition between DFG_in_ and DFG_out_ conformations—often induced by type-II inhibitors—this binary perspective does not fully capture the inherent conformational heterogeneity in apo kinases or how various perturbations shape their conformational ensembles. Our physics-based molecular simulations and Markov state models reveal that apo serine/threonine kinases (STKs) occupy a dynamic ensemble of three key macrostates—**DI_N_**, **DI_F_**_1_, and **DI_F_**_2_—whose populations can be fine-tuned by different conformational perturbations. Notably, shifting from DI_N_ to DI_F1_ and DI_F2_, where the DFG–Phe points toward the αC-helix, increases the fraction of broken Lys–Glu interactions, ultimately driving the αC-helix from its “*in*” to “*out*” conformation. Transitioning to DI_F1/F2_ state together with αC-helix “out” conformation opens up allosteric backpocket which can be targeted by type-III small molecules to develop selective therapeutics against kinases^37,40^.

Our simulations reveal that the V600E mutation in BRAF promotes a shift from the DI_F1_ to the DI_N_ population, activating the MAPK pathway and driving cancer cell proliferation. Vemurafenib, which binds to the hydrophobic back pocket of V600E BRAF, restores and stabilizes the DI_F1_ population. Likewise, other V600E-specific BRAF inhibitors—such as Dabrafenib and Encorafenib^41^—also stabilize the DI_F1_ state. Notably, our computational approach predicts that maintaining BRAF and MEK1 in the DI_F1_ conformation can restore normal cell division in V600E mutant cells. Building on this insight, we propose designing dual-target small molecules that bind the active sites of both BRAF and MEK1, shifting the population from DI_N_ to DI_F1_—an approach that could serve as a combination therapy for treating V600E-driven cancers.

Beyond active-site inhibitors, designing novel classes of therapeutics offers an attractive strategy to reduce off-target toxicity and combat resistance. By capturing the DI_F1_ conformational ensemble in the V600E BRAF–MEK1 complex, we identified an expanded hydrophobic backpocket in MEK1. Small-molecule “glues” (such as NST-628) can target this site to stabilize the V600E BRAF–MEK1 complex in the DI_F1_ state, thereby restoring normal cellular function^30^. The ability to target an expanded hydrophobic backpocket in MEK1 and stabilize RAF and MEK1 in DI_F1_ state is an attractive strategy to develop pan-RAF-MEK1 molecular glue degraders.

Beyond small molecule binding selectivity, the ability to stabilize distinct conformational states is crucial for determining protein–protein interactions, which in turn govern key cellular functions and downstream signaling. For example, the E3 ligase XIAP binds to RIPK2 and stabilizes the D_IN_ state (PDB: 8AZA), whereas the interaction between BRAF and MEK1 selectively stabilizes the DI_F1_ state in normal cells. Small-molecule inhibitors of RIPK2, such as SB-203580 (PDB: 5AR4^42^), and CSLP18^43^ (PDB: 6FU5), stabilize the DI_F1_ state, whereas molecules like K252 (PDB: 8X2O) favor the DI_N_ state. These observations suggest that DI_F1_ stabilizers act as PPI inhibitors, while DI_N_ stabilizers promote PPI interactions. Small molecules that stabilize the DI_F1_ conformation shift the equilibrium from the AC_out_ to the AC_in_ state, triggering a transition of the activation loop from an extended to a helical (collapsed) conformation (Figure S12 in Supporting Information). In the extended state, Arg171 forms a critical interaction with the E3 ligase XIAP. This extended-to-collapsed transition, along with an allosteric flip in Ile208 (**Figure 9**), modulates RIPK2–XIAP complex formation^21^. Because XIAP recruitment by RIPK2 is a key structural event in the NOD1/2 signaling pathway, our findings suggest that small molecules favoring the DI_F1_ state will exhibit excellent selectivity for RIPK2 and inhibit the RIPK2–XIAP interaction, thereby blocking NOD1/2 signaling.

This highlights the importance of understanding conformational heterogeneity and population shift in kinases to gain mechanistic insights into protein-protein interactions which modulate downstream signaling pathways implicated in inflammatory diseases. Our future study will combine machine learning augmented adaptive sampling and biochemical experiments to systematically characterize the effect of small molecules as on RIPK2-XIAP interactions, providing critical mechanistic insights into how small molecule binding allosterically modulates protein-protein interactions.

Recently, an enhanced sampling approach combining MSA subsampling, molecular dynamics (MD), and metadynamics with a variational autoencoder was applied to investigate the conformational transitions between DFG_in_ and DFG_out_ states^44^. In a follow up study, the authors applied this approach to kinase inhibitors^45^, demonstrating how conformations selected from the DFG_in_ and DFG_out_ ensembles can be used for molecular docking. However, both these studies fail to capture how ligand binding modulates the conformational dynamics^46^ within the DFG_in_ ensemble and allosterically modulates protein–protein interactions which governs downstream cellular signaling.

Here, we introduce a novel ML-augmented adaptive sampling protocol designed for two main purposes: (a) to capture functionally relevant slow conformational motions in different protein domains (**Figure S6** and **S13** in Supporting Information), revealing conformational allostery, and (b) to enable improved structural ensemble prediction through clustering in the latent layer of the ML model. In this scheme, short unbiased MD simulations are launched from the ensembles generated in the latent space, which, when combined with underlying physics, capture a more complete spectrum of conformational heterogeneity in both apo and holo kinases. This approach goes beyond what can be achieved by AlphaFold-seeded simulations or traditional ultra-long MD.

A key feature of our protocol is the latent layer’s ability to provide a compressed, two-dimensional representation of structural features originating from distinct high-dimensional inputs, such as the N-lobe and C-lobe dynamics of RIPK2. This separation of structural features is crucial for capturing slow conformational motions in larger proteins. We further demonstrate that our ML model captures the DI_N_ to DI_F1_ transition in RIPK2 and show how this transition allosterically influences the XIAP binding interface that governs RIPK2–XIAP complex formation. Additionally, we have highlighted how the latent layer of the ML model can capture ‘*slow*’ conformational dynamics of the BRAF-MEK1 complex upon V600E mutation (**Figure S3** in Supporting Information). Finally, we highlight how the latent representations derived from the ML model can be employed as collective variables^47^ for constructing MSMs (**Figure S9** and **S10** in Supporting Information) and for driving enhanced sampling simulations such as metadynamics.

Understanding how small-molecule binding modulates protein dynamics^48^ without altering the underlying potential energy surface remains a major challenge, as conventional molecular simulations have limited ability to capture slow timescale conformational changes and sampled rugged free energy surface. Here, starting from the Vemurafenib-bound DI_N_ state of V600E BRAF, we demonstrate how one can sample **DI_N_**-to-**DI_F1/F2_** conformational transitions using a multi-step protocol: (a) launch multiple independent short MD simulations, (b) train a machine learning (ML) model and generate conformational ensemble from its latent layer, (c) initiate new MD simulations from these latent-layer conformations, and (d) combine all simulations into a Markov state model (MSM). This workflow captures both the population shift and conformational heterogeneity of the holo V600E BRAF complex. Traditional structure-based drug discovery methods often rely on ranking small molecules solely by free energy values. However, consider a scenario where several compounds bind to a kinase with nearly identical free energy values, yet produce distinctly different effects on kinase-mediated protein–protein interactions. Conventional free energy approaches (e.g., FEP^49^ or MM/GBSA^50^) struggle to capture these differences because they do not account for shifts in the conformational ensembles upon ligand binding^46^. Our adaptive sampling protocol directly addresses this limitation by revealing how small molecule binding modulates protein’s conformational states. By illuminating transitions between distinct conformational states, this method explains how kinase-mediated protein–protein interactions can be modulated in different ways by ligands with similar free energy profiles. More broadly, the approach is applicable to any protein–ligand system and provides critical insights for designing highly selective therapeutics capable of differentially modulating protein–protein interactions through fine-tuning of protein dynamics.

Our ensemble definition extends beyond the **DI_N/F1/F2_**states to include the conformational heterogeneity of the activation loop, as exemplified by CDK2. Although the DFG-Phe of CDK2 remains in the **DI_N_**conformation, the activation loop remains in a dynamic equilibrium between collapsed (**AC_in_**) and extended (**AC_out_**) states. The activation loop dynamics, in turn, modulate the **LG_L/U_**conformations. Understanding this interplay—particularly how the activation loop dynamics and the Lys-Glu interaction influence small-molecule binding or protein–protein interactions—is crucial for elucidating CDK2 signaling pathways. For instance, binding of selective small molecules to CDK2^51^ (PDB: 8UV0, 9FR2, 7ZPC, 7SA0, 7S9X) shifts the conformational equilibrium toward the **AC_in_LG_U_**state, whereas CyclinE1 binding stabilizes the **AC_out_LG_U/L_** ensemble. Recognizing that CDK2’s conformational dynamics extend beyond DFG-Phe is key to designing selective small molecules that induce the **AC_in_** conformation or developing molecular glues that exploit the cryptic pocket at the CDK2–CyclinE1 interface. We propose that the conformational heterogeneity in CDKs is foundational to their interactions with cyclins and is key to develop selective therapeutics modulating specific CDK-Cyclin interactions.

Our ensemble definition can be extended to capture conformational heterogeneity in the DFG_out_ ensemble (**Figure 15, 16** and **Figure S16**, **S17** in Supporting Information) and is not limited to serine/threonine kinases. Notably, our adaptive sampling framework prospectively sampled **DI_N to_ DI_F1_** conformational transitions in the tyrosine kinase FGFR2 that was not detected by ultralong simulations by DE Shaw Research^40^ initiated from the apo structure or traditional AlphaFold seeded molecular simulations (**Figure S15** in Supporting Information). Our approach, leveraging AlphaFold-seeded molecular simulations, successfully captured the full spectrum of conformational heterogeneity in apo ABL kinase, a crucial drug target for chronic myeloid leukemia. While previous studies required a 10-millisecond ultra-long MD simulation to sample transitions between the DFG_in_ and DFG_out_ conformational states, our protocol achieved this 80- fold faster (total simulation length 8 microsecond). Beyond accelerated sampling, our workflow also uncovered novel conformational states that have remained elusive due to insufficient sampling and a lack of clear ensemble definitions. Although NMR experiments have revealed some conformational heterogeneity in apo ABL kinase, they did not identify two additional DFG states that our computational method discovered (**Figure 15**). We believe this workflow offers a significant advancement, enabling *a priori* conformational sampling in biomolecular simulations with a fraction of computational cost compared to ultralong MD simulations.

**Figure 15.**
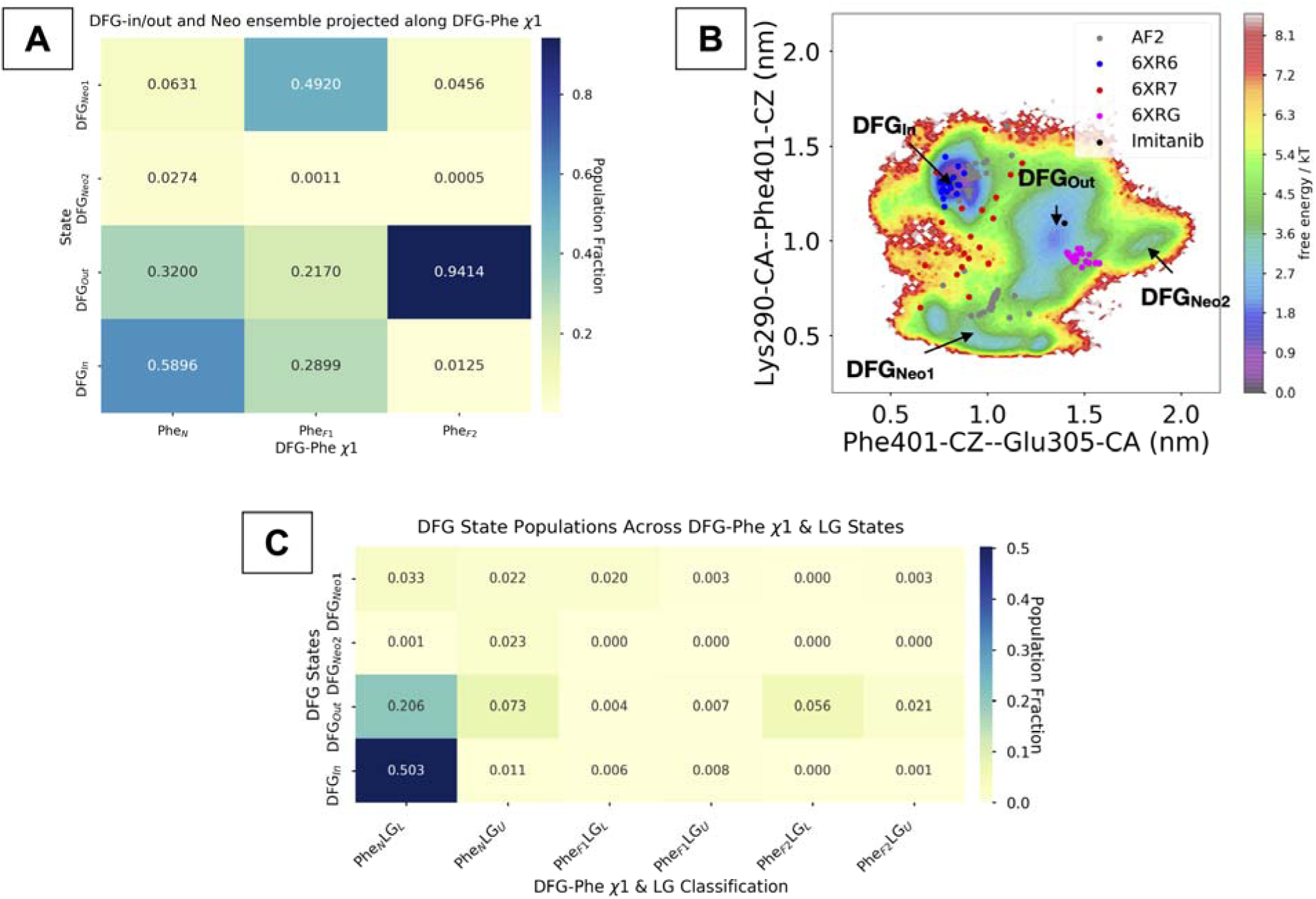
**A**: We characterized the conformational heterogeneity of apo ABL kinase by determining the MSM-weighted populations of DFG_in/out/Neo1/Neo2_ substates, using Phe_N_, Phe_F1_, and Phe_F2_ as key descriptors. **B**: An MSM-weighted free energy surface, projected along two crucial distances, highlights the superior sampling achieved by AI-augmented molecular simulations compared to both NMR (PDB: 6XR6, 6XR7, 6XRG) and AlphaFold2-generated conformational ensembles. Notably, our simulations also successfully captured holo-like structures of ABL kinase, consistent with the imatinib-bound state. **C**: The conformational heterogeneity across DFG_in/out/Neo1/Neo2_ substates was further elucidated through ensemble definitions based on the DFG-Phe χ1 angle and the Lys290-Glu305 salt bridge interactions (LG_U/L_). The population weight was calculated using *sin* and *cos* transformed DFG-Phe χ1/χ2 angles. The convergence of MSM weighted populations across different lag times is highlighted in Figure S18 in Supporting Information.

**Figure 16.**
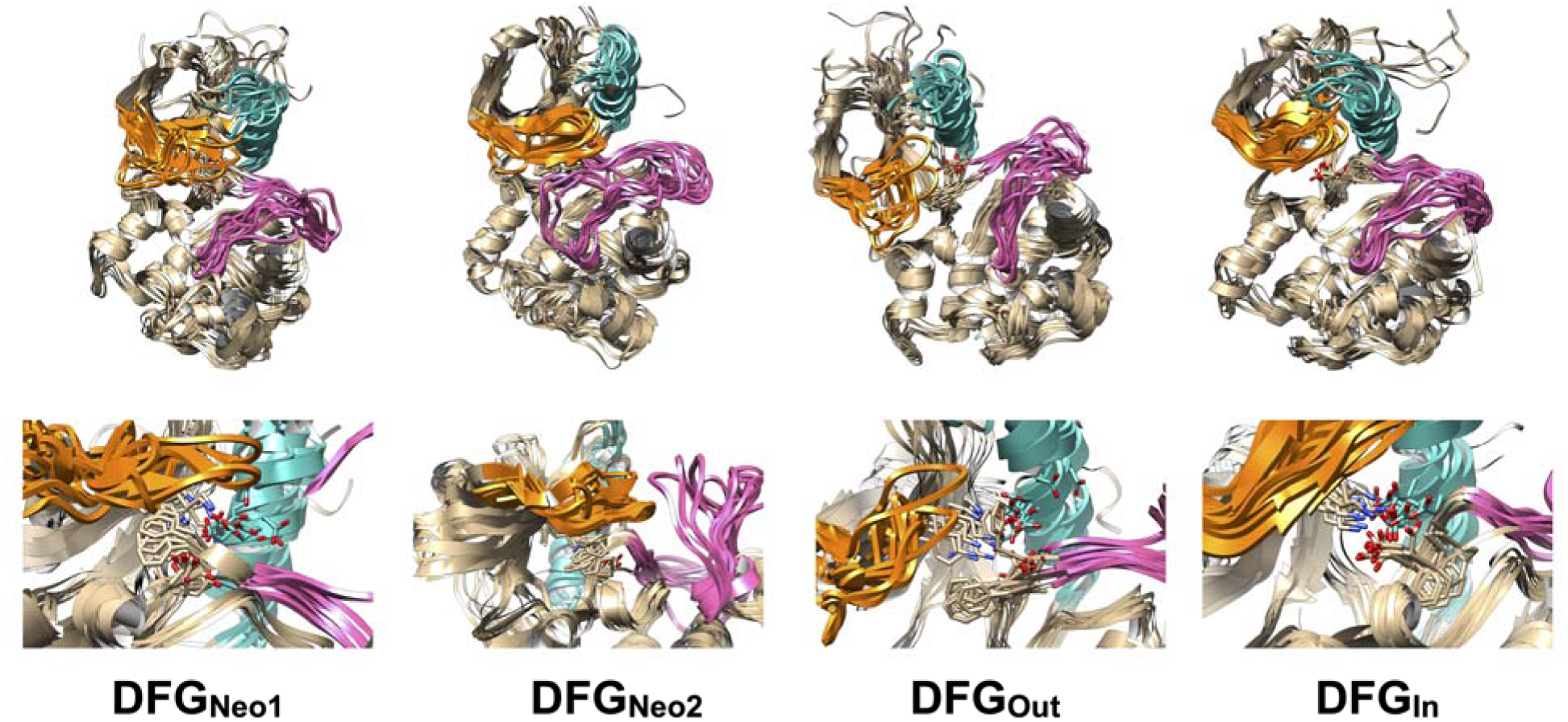
The upper panel illustrates the conformational ensemble of ABL kinase, highlighting structural heterogeneity in its P-loop (orange), *αC-helix* (light sea green), and activation loop (pink) across different DFG sub-states. The lower panel provides a closer view, showing the local conformational heterogeneity of DFG-Asp, DFG-Phe, Lys290, and Glu305 (stick representation) within these DFG sub-states.

Some may argue that the state populations we report merely reflect the choice of established collective variables (CVs). Yet for ABL kinase we showed that a machine-learning model trained only on dihedral angles of two distinct kinase domains recovers the same underlying free-energy landscape as expert-defined CVs. While absolute state populations vary, the shape of the free energy landscape and the shifts among metastable states remain consistent (**Figure S19** in Supporting Information). This agreement highlights the robustness of our accelerated sampling approach for revealing functionally relevant ABL kinase conformations. In future, computational ensemble generated from our protocol can be augmented with NMR-derived conformational ensembles, providing a more comprehensive understanding of protein dynamics and conformational heterogeneity.

The ability to prospectively sample conformational heterogeneity in kinases will be pivotal to integrate our accelerated computational pipeline with downstream traditional or AI augmented^52,53^ population shift^54^ based virtual screening pipeline to identify and develop selective therapeutics targeting a wide range of kinases. Furthermore, the ability to distinguish conformational populations across kinase homologs, as demonstrated for IRAK1/4 and FGFR1/2, can be generalized to other kinase families. This approach will enable the design of homologue-specific therapeutics with reduced off-target effects.

## Conclusion

Our study underscores the importance of human interpretable ensemble definitions that capture not only endpoint structures but also the populations and timescales associated with conformational heterogeneity in protein subfamilies. Much of the current work in generative AI for protein structure prediction focuses on sampling rigidly defined conformational endpoints^13,14,55^, overlooking the continuous spectrum of functionally relevant states and limiting our ability to understand how perturbations shift populations and timescales. The accelerated molecular simulation frameworks introduced here, combined with human interpretable ensemble definitions, can be extended to other protein subclasses—such as GPCRs, tumor necrosis factors (TNFs), intrinsically disordered proteins (IDPs)^56^ —where conformational heterogeneity drives molecular recognition^57^.

A major bottleneck to capturing rare yet functionally important conformational states—and thereby learning a generalized model of protein dynamics with interpretable collective variables—is the scarcity of training data. Our work emphasizes that high-quality conformational dynamics data is crucial for developing a generalizable understanding of protein dynamics. The recently proposed deep learning framework, *BioEmu*^58^, has garnered attention for suggesting it may substitute molecular dynamics (MD) simulations. However, *BioEmu* cannot predict the thermodynamics and kinetics of population shifts under ensemble perturbations, nor is it capable of capturing generalized population shifts across different protein classes—primarily due to insufficient training data and inability to capture of ‘*out of distribution*’ populations in the absence potential energy functions. In contrast, the computational framework described in our study can overcome these limitations by accurately modeling rare and functionally relevant conformational ensembles of biomolecules. The sampling approaches proposed in this study, combined with the diverse conformations available in the Protein Data Bank, can create high quality datasets needed to develop protein ensemble predictors from a given sequence within a specific protein subfamily. In the future, ensemble generation from ML model’s latent layers, integrated with flow matching^59^/diffusion models^60^ and Langevin dynamics^61^, will provide a pathway to capture the full breadth of conformational heterogeneity across protein subfamilies—without requiring extensive molecular dynamics or enhanced sampling.

## Methods

### Structural ensemble generation using AlphaFold

A structural ensemble for each sequence of serine-threonine kinases (wild/K47R RIPK2, monomeric wild/V600E BRAF, CDK2, IRAK1, IRAK4, FGFR2, MAPK14, ABL) was generated using the ColabFold^62^ implementation of AlphaFold2 (link: https://colab.research.google.com/github/sokrypton/ColabFold/blob/main/AlphaFold2.ipynb#scrollTo=ADDuaolKmjGW), following the protocol outlined by Meller and co-workers^12^. We first built an initial multiple sequence alignment (MSA) via the MMseqs2^63^ method integrated in ColabFold. Next, the MSA was subsampled stochastically to include up to 16 cluster centers and 32 additional sequences (referred to as *max_msa* = 16:32). For ensemble generation, we used the “*greedy*” pairing strategy to match any taxonomically compatible subsets. We used 16 random seeds and enabled model dropout. Leveraging both model dropout and multiple seeds capitalizes on AlphaFold2’s^64^ model uncertainties, resulting in 80 predicted structures per protein and capturing conformational heterogeneity.

Five protein–protein complexes, involving wild-type and V600E BRAF–MEK1 as well as CDK2–CyclinE1, were generated using AlphaFold3^65^ (https://alphafoldserver.com/). The objective was to assess whether AlphaFold3 can capture the full spectrum of population shifts and to determine if molecular simulations initiated from these AlphaFold3-generated conformations can reveal conformational heterogeneity.

### Molecular dynamics simulations

Each apo protein structure was prepared for molecular dynamics simulations using the *tleap* module from Amber2022^66,67^, following the protocols outlined by Meller et al.^12^ and Vats et al.^21^ Proteins were parameterized with the AMBER FF14SB^68^ force field. For holo simulations, the small molecule (ligand) was parameterized with the GAFF^69^ force field and partial charges were added using AM1-BCC^70^ method. Counterions were added to each system, and the systems were then solvated in a truncated octahedron box containing TIP3P^71^ waters, ensuring a minimum distance of 10 Å from the protein to the box boundary.

A two-phase energy minimization was performed on each system: a) Phase 1: Only the solvent and ions were minimized while restraining the protein with a 100 kcal/mol·Å² potential (200 steps of steepest descent followed by 200 steps of conjugate gradient), b) Phase 2: The entire system was minimized without restraints for 500 steps conjugate gradient.

Following minimization in Amber2022, the Amber topologies were converted to Gromacs format using Acpype^72^. Each system was then gradually heated from 0 K to 300 K over 500 ps in the NVT ensemble, applying harmonic restraints (500 kJ/mol·nm²) on the backbone of heavy atoms. Subsequently, the restraints were removed, and the systems were equilibrated for 200 ps in the NPT ensemble at 300 K, with pressure maintained at 1 bar by the Parrinello–Rahman barostat^73^ and temperature controlled by the v-rescale thermostat.

Production simulations were conducted in the NPT ensemble at 300 K and 1 bar using the leapfrog integrator with a 2 fs timestep. A 1.0 nm cutoff was used for nonbonded interactions, while long-range electrostatic interactions were treated via the Particle Mesh Ewald (PME)^74^ method with a 0.16 nm grid spacing. The LINCS^75^ algorithm constrained covalent bonds to hydrogen atoms. All heating, equilibration, and production runs were carried out using GROMACS 2022^76^.

For the AlphaFold-seeded molecular dynamics of monomeric RIPK2, CDK2, BRAF (wild and V600E), FGFR2, IRAK1/4, and MAPK14, a total of 80 structures were generated by AlphaFold. Each structure underwent a 100 ns simulation, culminating in 8 μs of total simulation time (80 × 100 ns).

Wild/V600E BRAF–MEK1 and CDK2-CylcinE1 complexes were generated using AlphaFold3 and prepared according to the aforementioned protocol. For each complex, five structural predictions were generated, and two independent 200 ns simulations (5 snapshots * 2 replicas * 200 ns each = 2000 ns) were performed for each prediction. Additionally, RIPK2+XIAP simulations from *Vats et al*.^21^ were used for post processing. In all cases, trajectories were saved every 10 ps.

### Slow Feature Analysis

Slow Feature Analysis (SFA)^77^ is a dimensionality reduction technique designed for high-dimensional temporal data. Its primary objective is to transform the *J*-dimensional input signal, ▢(▢), into output signals ▢_▢_(▢) = ▢_▢_(▢(▢)) using a set of nonlinear functions gk(c). These output signals are optimized to minimize

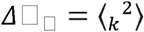

where _k_ represents the temporal derivative of ▢_▢_(▢), and ⍰⍰⍰ denotes temporal averaging. By minimizing *Δ*▢_▢_, SFA extracts features that vary slowly over time.

Vats et al.^21^ highlighted how SFA can capture linear combinations of slowly varying structural features from high-dimensional data generated from molecular dynamics (MD) simulations. In this study, we demonstrate how SFA can be applied to capture slowly varying conformational dynamics in kinases. Furthermore, we utilized SFA to develop the Markov state model^15,16^ (Figure S10 in Supporting Information), which enabled us to predict the populations of functionally relevant conformations in kinases.

### Time-lagged autoencoder

A time-lagged autoencoder (TAE)^19^ is a specialized neural network architecture designed to learn compact representations of time-series data while explicitly leveraging temporal structure. This approach is particularly powerful in scenarios such as molecular dynamics (MD) simulations, where the goal is to capture slowly varying conformational dynamics of biomolecules from high-dimensional data. Identifying these long-timescale conformational transitions is crucial for understanding phenomena like conformational allostery in kinases.

Unlike standard autoencoders—which aim to reconstruct the same input data—time-lagged autoencoders are trained to predict *x_t+τ_* from *x_t_*, where τ is a user-defined lag time. By focusing on a non-zero *τ*, the TAE learns latent representations *z_t_* that capture the slowly evolving features of the molecular system, which are relevant for predicting future states.

The TAE architecture comprises a) Encoder: Transforms the current system state *x_t_* into a lower-dimensional representation *z_t_*. b) Decoder: Reconstructs the time-lagged version of the system, *x_t+τ_*, from the latent representation *z_t_*.

By pairing *x_t_* and *x_t+τ_*, the model naturally emphasizes slowly varying dynamics, since the most prominent changes occurring over lag time τ are embedded in the latent space.

Latent Layer Representation: Let *x_t_* ∈ ℝ*^d^* be the high-dimensional molecular state at time t. In the encoder, the latent layer is often formulated as:

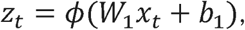

where:

● *W*_1_ ∈ ℝ^(m×d)^ is the encoder’s weight matrix.
● *b*_1_ ∈ ℝ*^m^* is the bias vector.
● *ϕ*(.) is a nonlinear activation function.
● *z_t_* ∈ ℝ*^m^* is the low-dimensional latent representation.

The decoder then reconstructs *x_t+τ_* from *z_t_*, typically using:

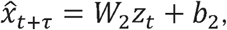

Where *W*_2_ ∈ ℝ^(d×m)^ and *b*_2_ ∈ ℝ*^d^*. The training objective is to minimize the reconstruction error:

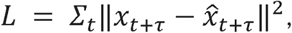

which encourages *z_t_* to preserve information about the slow degrees of freedom predictive of future conformations.

The TAE’s learned latent space provides a compressed, nonlinear representation of input features that captures slowly varying conformational dynamics in molecular systems. Our machine learning framework adopts a dynamic feature-selection strategy by applying a learnable masked kernel through a ReLU activation, ensuring that only features with non-zero weights are retained. This yields two clear benefits: the retained features are easier to interpret, and they can be more effectively used as collective variables (CVs) in enhanced sampling simulations.

To give the latent space more structure, we extended the method with two modules: (a) We infused Gaussian noise into the output of the encoder before further processing it with the decoder, which as a result enforces better separated long lived states. (b) we expanded the training objective with the VAMP score^78^ estimated in the latent space, thereby ordering the long-lived states by the slowest motions.

During training, the “*CVEstimator*” module orchestrates the entire process by balancing the reconstruction loss, the penalty for large, masked kernel weights, and the VAMP loss—each weighted appropriately—to optimize both interpretability and dynamical fidelity. This integrated approach results in an end-to-end pipeline where feature selection, sparse representation learning, and dynamical analysis via VAMP collectively yield a robust and physically meaningful representation of the input data.

This setup enabled us to identify slowly evolving structural features in various protein domains. These nonlinear combinations of structural features serve as collective variables for constructing Markov state models (**Figure S4** in Supporting Information) or can be incorporated into enhanced sampling frameworks. In our study, we used sine/cosine-transformed dihedral angles of different kinase domains as inputs, with two latent layers to capture compressed representations of each feature set.

### Adaptive sampling

Our adaptive sampling method preserves the underlying dynamics by restarting MD trajectories at carefully chosen seeds, without introducing biasing forces. This approach addresses the sampling challenges that can arise from AlphaFold-seeded molecular dynamics simulations. The sampling pipeline (**Figure 17**) follows these steps:

**1. Starting conformations: Apo Simulations:** Short (10 ns each) independent molecular dynamics (MD) simulations were launched from an AlphaFold-generated conformational ensemble (total 80 structures, training data: 10 ns * 80 = 800 ns) of apo IRAK1 and FGFR2 obtained through MSA subsampling. **Holo Simulations:** Small molecule was placed in the active site of the DIN conformational state of V600E BRAF. From this holo starting structure, 20 independent 10 ns MD simulations (10 ns * 20 = 200 ns) were initiated.
**2. Time-Lagged Autoencoder:** A time-lagged autoencoder was trained on the sin/cos- transformed χ1 and χ2 angles of the αC-helix and the activation loop in kinases (apo IRAK1, FGFR2 and the holo V600E BRAF monomer). This captures slowly varying conformational features from high-dimensional MD trajectory data (**Figure S6** and **S13** in Supporting Information).
**3. Sampling conformations from the latent layers:** *K-center*^79^ clustering (K=100) was performed on the 2D latent space produced by the time-lagged autoencoder. Frames corresponding to each cluster center were extracted, representing the ’*physics-refined’* conformational ensemble.
**4. Refined Simulations and Markov State Model:** Independent 100 ns MD simulations were launched from each cluster center (total 100ns * 100 replicas = 10 microsecond) which were combined using the Markov state model to capture populations and timescales associated with conformational transitions.

**Figure 17.**
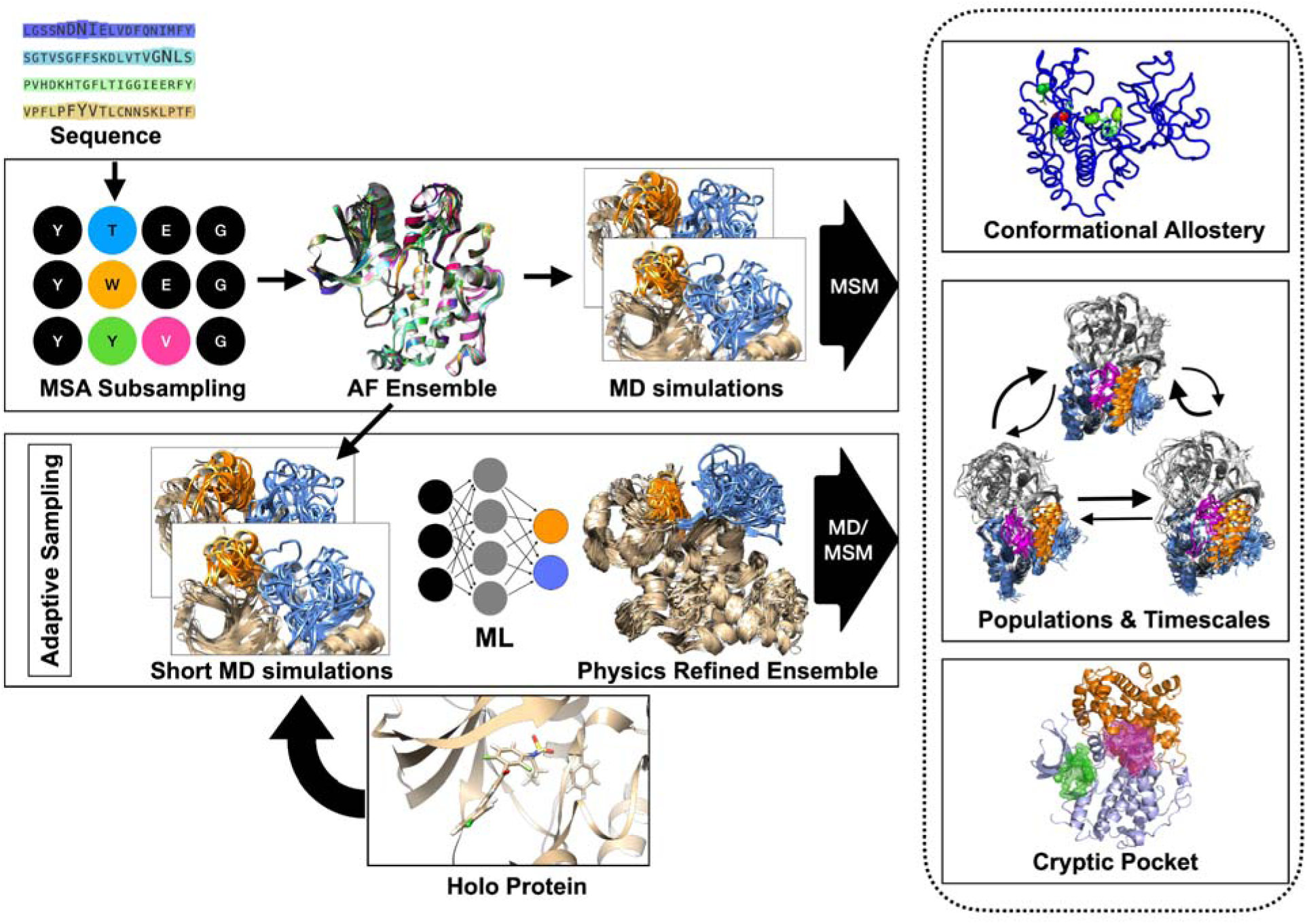
Overview of the computational framework employed in this study to capture generalizable dynamics in kinases and how perturbations such as ligand binding/mutations modulate conformational dynamics. The framework requires a protein sequence as input and uses the MSA subsampling approach to generate initial structural guesses (AF ensemble) using AlphaFold. These structures were then used to seed to launch independent MD simulations and finally combined together using the Markov state model (MSM) to capture conformational dynamics in kinases. In some cases, this standard workflow may not fully capture the breadth of conformational heterogeneity. To address such scenarios, we used an adaptive sampling framework which starts with launching short MD simulations (10 ns) from an AF generated conformational ensemble. The high-dimensional data from these simulations are used to train a time-lagged autoencoder (TAE), which identifies slowly varying structural features. Clustering in the TAE’ latent space yields a “physics-refined” ensemble, and representative cluster centers are selected as starting points for further MD simulations. This expanded ensemble is then integrated into the MSM, providing a more comprehensive view of kinase conformational heterogeneity in both apo and holo states. Capturing these kinase dynamics facilitates the study of conformational allostery and reveals druggable cryptic pockets in protein–protein complexes, potentially accelerating the development of targeted therapies.

### Markov state model

Markov state models (MSMs) are a powerful framework for analyzing the conformational dynamics of biomolecules, providing a probabilistic description of the transitions between discrete states in a high-dimensional conformational landscape. MSMs leverage time-resolved data, such as that from molecular dynamics simulations, to model the system as a network of states connected by transition probabilities, capturing the likelihood of moving from one state to another over a defined lag time. Key steps in constructing an MSM include defining the conformational states—commonly achieved through clustering methods like k-means—and estimating the transition probabilities using a maximum-likelihood approach. By evaluating the dynamics at different lag times, MSMs enable the extraction of equilibrium populations and characteristic timescales of conformational transitions.

Markov state modeling (MSM) was performed on the sine and cosine transformations of the χ1 and χ2 angles of DFG-Phe, extracted from unbiased molecular dynamics simulations (launched from an AlphaFold-generated conformational ensemble) and from adaptive sampling simulations. K-means clustering (k = 100) was performed in the transformed dihedral space, and maximum-likelihood MSMs were generated using the lag times highlighted in the **Supporting Information**. Equilibrium populations obtained from these MSMs were projected along our ensemble definition to capture conformational heterogeneity in serine-threonine kinases. The convergence of MSM was assessed by predicting populations of different conformational states at different lag times (**Figure S9-S11, S2 and S4** in Supporting Information). PCCA+^32^ was employed to define macrostates that distinguish conformational states associated with DFG-Phe. The lag time of 10 frames was used to build final MSMs unless explicitly mentioned.

For apo RIPK2, we used the 2D latent layer of a TAE and the first two slow features extracted from slow feature analysis, illustrating how one can build an MSM on a compressed representation. MSM generations were performed using PyEMMA 2.5.7^80^.

## Data Availability

Inference code and the automated pipeline can be accessed here: https://github.com/svats73/mdml. Due to large size of the trajectories, they are available for non-commercial use by contacting the corresponding author. The ABL kinase trajectory can be accessed here: https://osf.io/4v7db/

## Supporting Information

Free energy surfaces, populations of conformational states at different lag times and ML weights for different protein kinases are available in the Supporting Information.

## Acknowledgement

Computational simulations were performed using Tufts High Performance Computing Cluster and standalone Linux Clusters with RTX4090 graphics cards.

## Competing interests

Authors declare no conflict of interests. Soumendranath Bhakat and Shray Vats are cofounders of AlloTec Bio, Inc.

## Notes

### Summary of Updates

Added result for ABL kinase (new subsection "AI accelerated molecular simulation captures DFGin to DFGout transitions in apo tyrosine kinase") and reformatted the manuscript accordingly to accommodate new results, figures (Figure 15 and 16) and discussions.

