## Supporting Information V2 for "Generalizable Protein Dynamics in Kinases: Physics is the key"

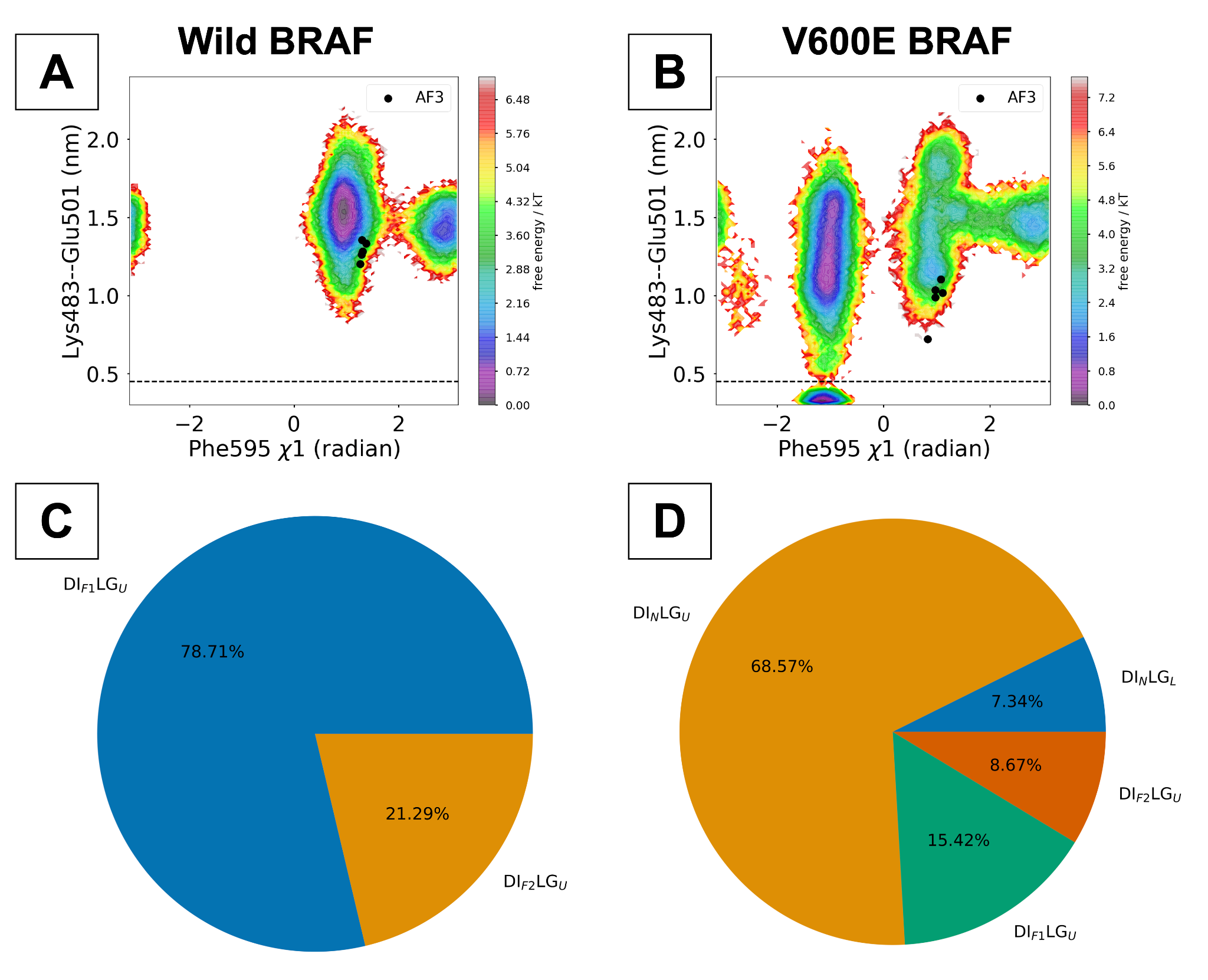

**Figure S1**. Free energy surfaces (A, B) and analyses of conformational heterogeneity (C, D) in wild-type and V600E BRAF–MEK1 complexes reveal that the V600E mutation shifts the conformational population from DI_F1_LG_L/U_ to DI_N_LG_L/U_ states, thereby activating the MAPK pathway and driving cancer cell proliferation.

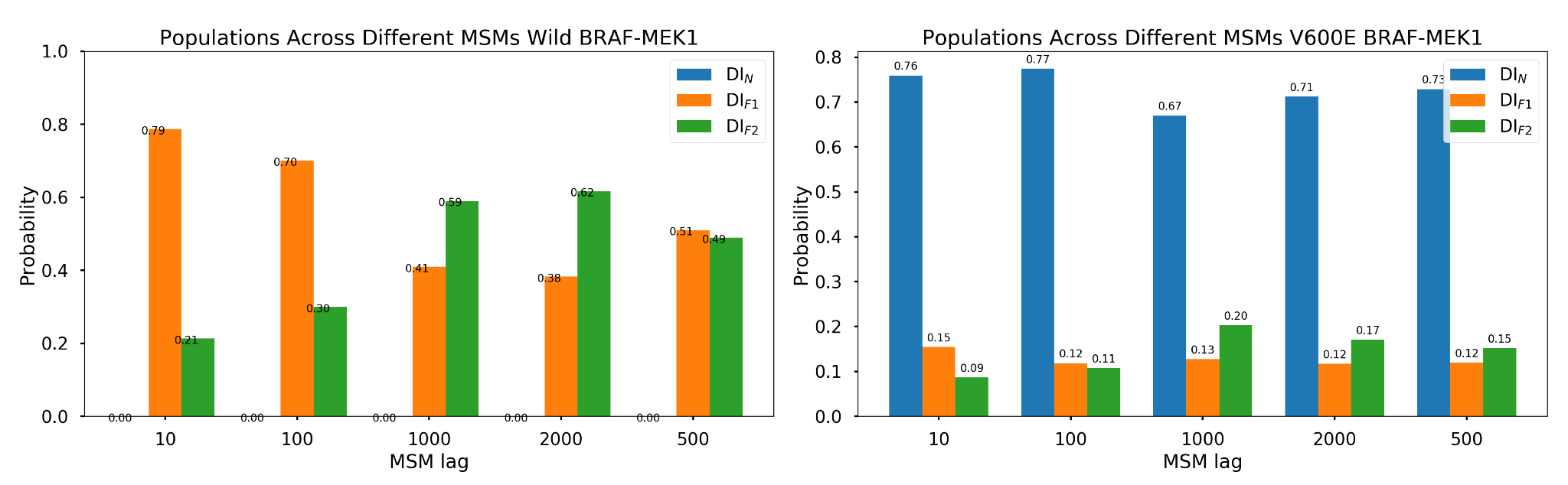

**Figure S2**. Probability of DI_N/F1/F2_ states as a function of lag time (in frames) for BRAF-MEK1 complexes. MSM was performed on sin/cos-transformed χ₁/χ₂ angles of DFG-Phe of BRAF. The population shift from DI_F1/F2_ to DI_N_ upon V600E mutation is consistent across the choice of lag times.

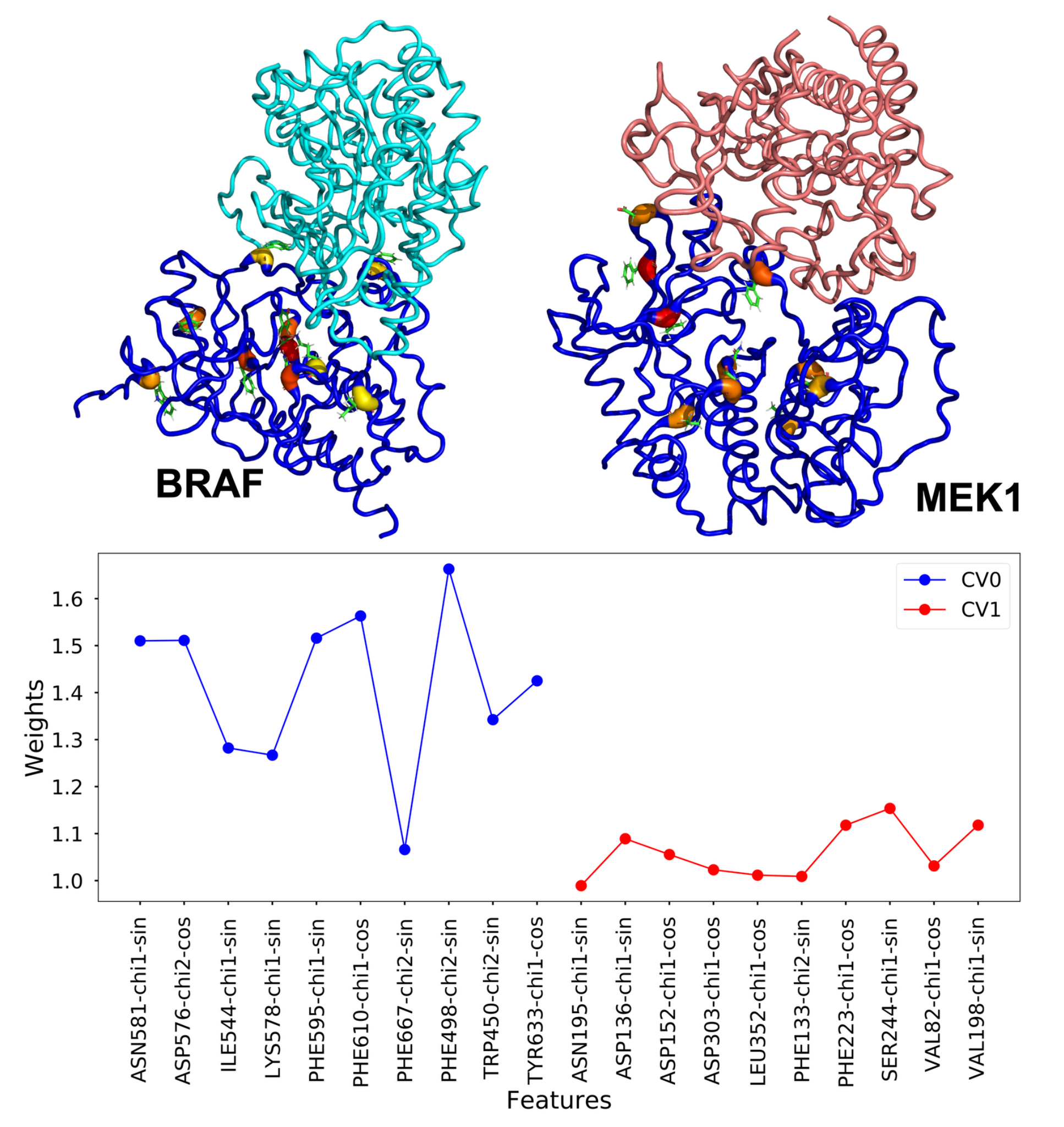

**Figure S3**. Latent layers of ML capture slowly varying conformational features in the BRAF (CV0) and MEK1 (CV1) subdomains of the V600E BRAF–MEK1 complex, illustrating how these latent layers can track protein domain dynamics. The corresponding ML weights are mapped onto the PDB structure and visualized as B-factors.

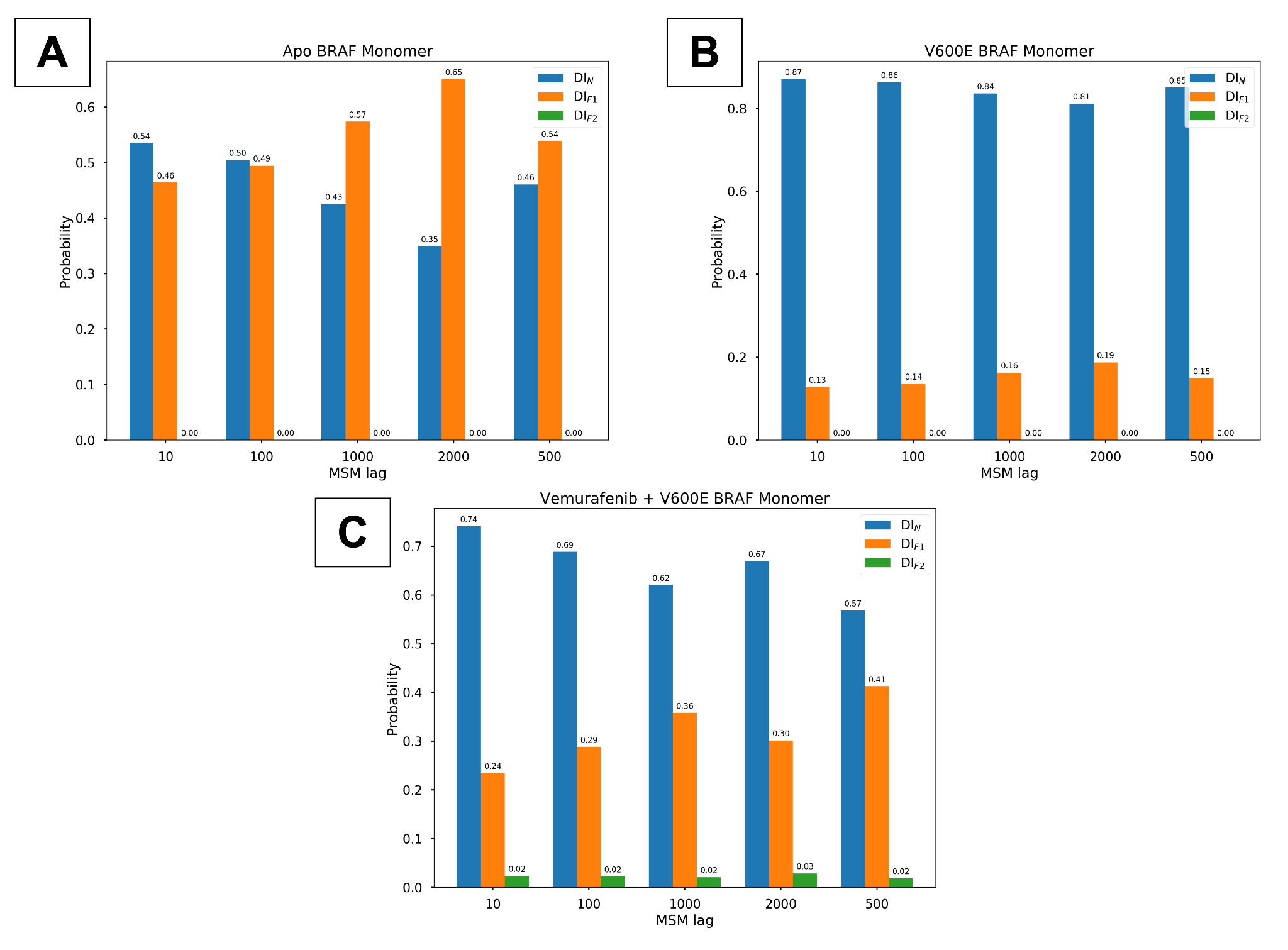

**Figure S4**. The population of DI_N/F1/F2_ conformational states as a function of Markov state model (MSM) lag times (in frames) captures the relative populations in the apo BRAF monomer and the population shifts induced by the V600E mutation and ligand binding. Consistent shifts across a range of MSM lag times underscore the convergence of these simulations. MSM was performed on sin/cos-transformed χ₁/χ₂ angles of DFG-Phe of BRAF.

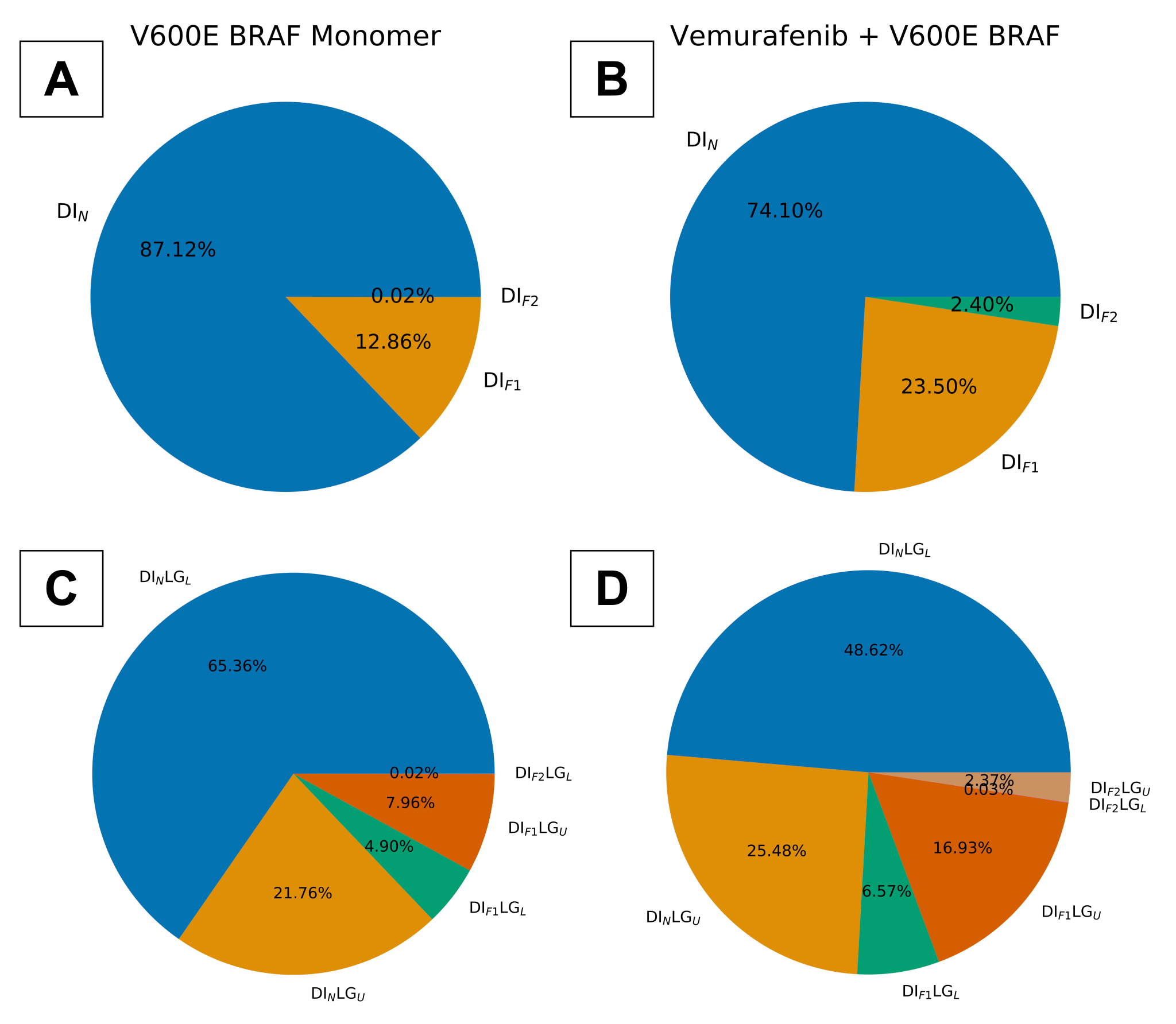

**Figure S5**. Population of conformational states in the V600E BRAF monomer and the vemurafenib-bound V600E BRAF monomer reveals how small-molecule binding shifts the population toward the DI_F1/F2_ states, as captured by our ensemble definitions. Population is predicted using sin/cos transformed dihedral angles of DFG-Phe and a lag time of 10 frames.

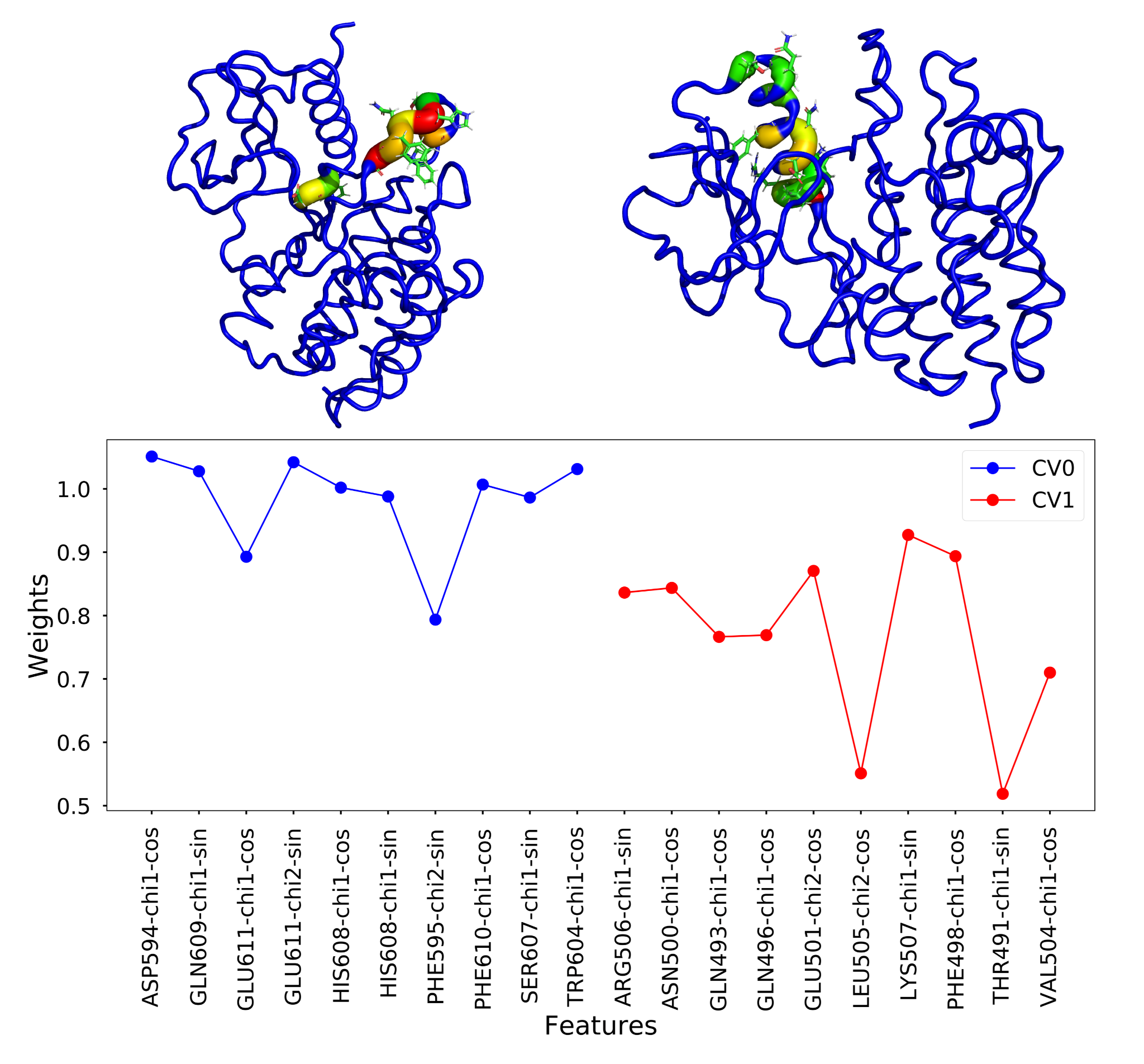

**Figure S6**. ML weights projected onto two latent layers—CV0, which captures the DFG and activation loop dynamics, and CV1, which captures the αC helix dynamics—reveal slowly varying structural features in 20 independent 10 ns MD simulations (total *200 ns*) initiated from the V600E BRAF DIN conformation. Simulations launched from physics-refined conformational ensembles derived from these ML latent layers explore the full spectrum of conformational heterogeneity in the holo V600E BRAF. ML weights are mapped to the PDB structure and highlighted as B-factors allowing for visualization of the key structural features captured by ML.

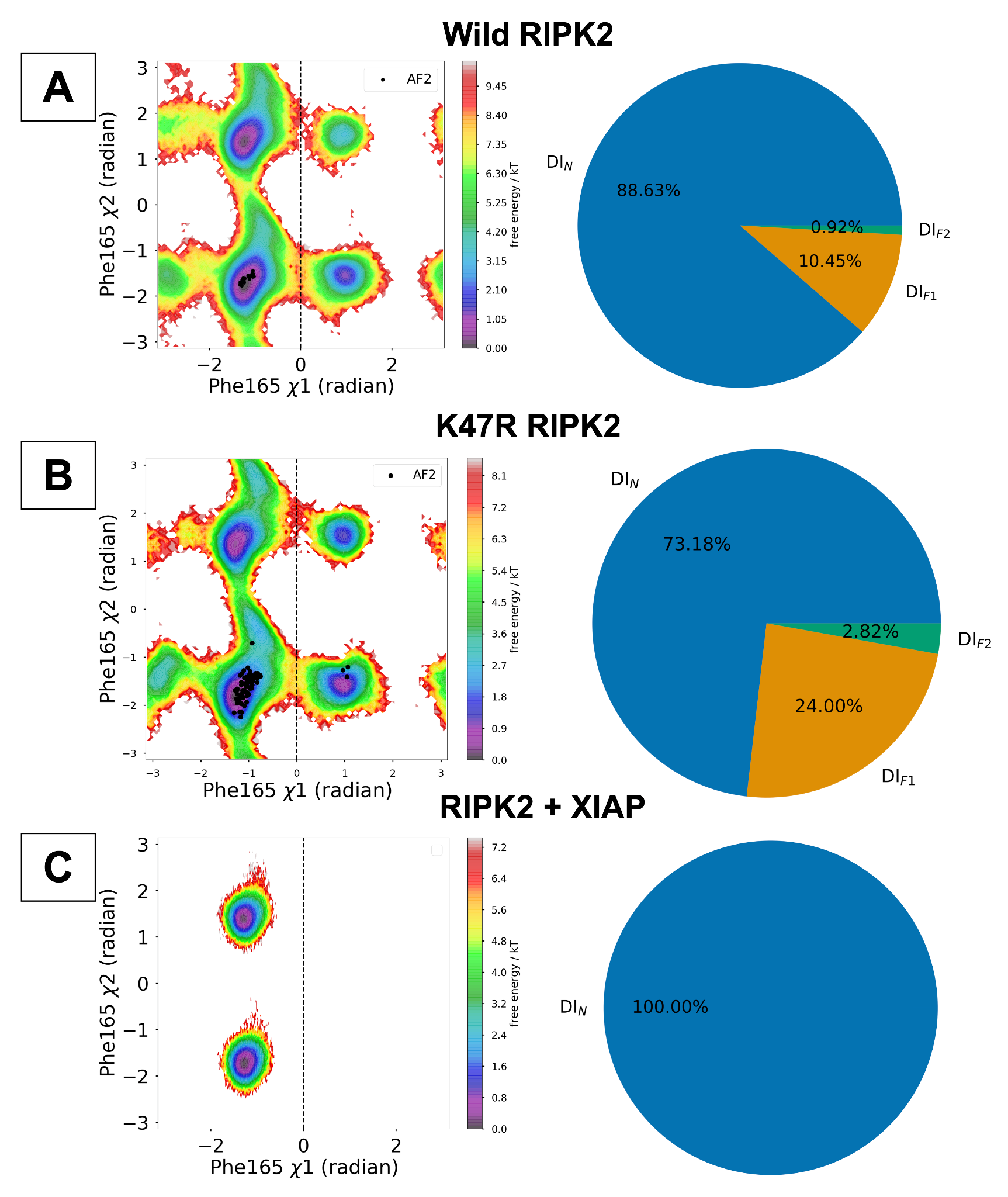

**Figure S7**. Free energy surfaces projected along the χ₁ and χ₂ angles reveal how different perturbations—such as the K47R mutation and interaction with the E3 ligase XIAP—alter the conformational populations of apo wild RIPK2 (A). The K47R mutation shifts the equilibrium from the DI_N_ state toward DI_F1_ (B), whereas XIAP binding stabilizes the DI_N_ state (C). It is critical to highlight that protein structure prediction models such as AlphaFold2 (highlighted as AF2) only provide static representation of protein but fail to predict how perturbations modulate protein dynamics.

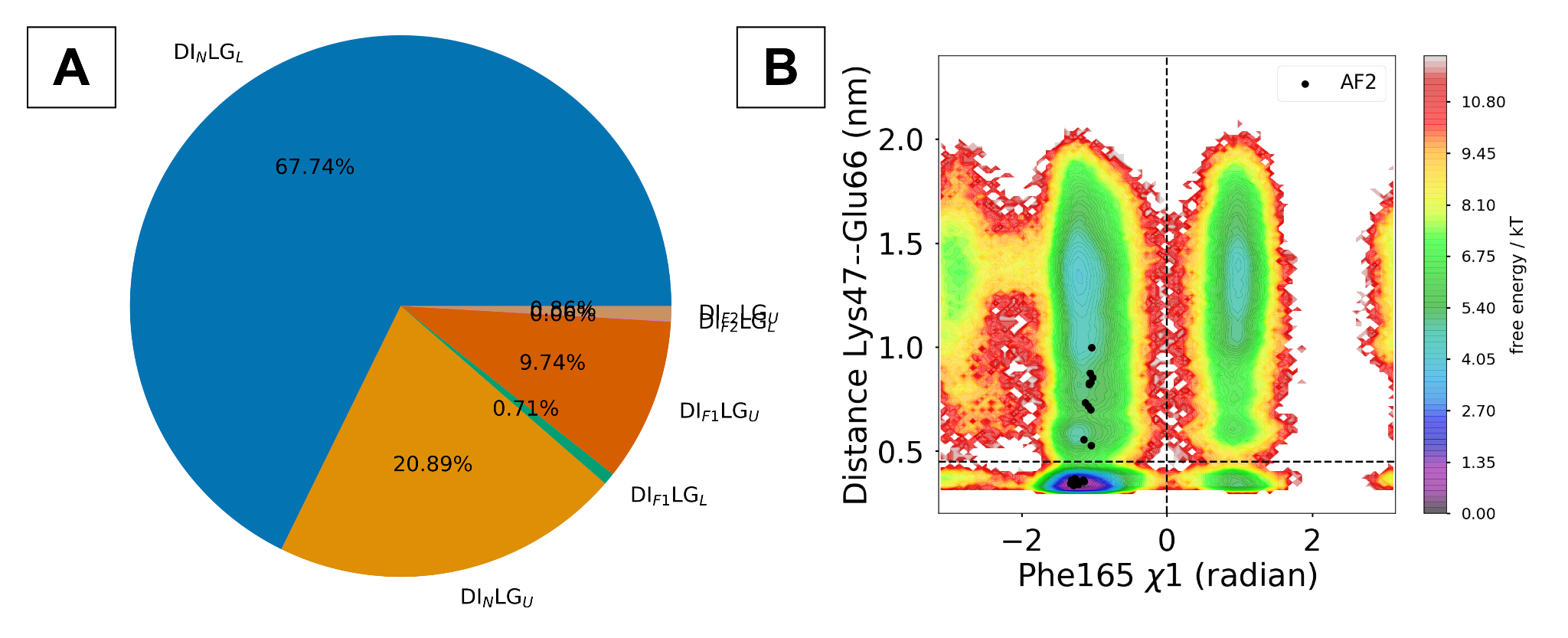

**Figure S8**. Populations (A) and the free energy surface (B) in apo wild-type RIPK2 demonstrate how transitions among DI_N_, DI_F1_, and DI_F2_ states capture the conformational heterogeneity between Lys–Glu-latched and unlatched conformations. In contrast, AlphaFold2 fails to capture both the transitions from DI_N_ to DI_F1_/DI_F2_ and the conformational heterogeneity within DI_F1_/DI_F2_ states (**DI_F1/F2_LG_L/U_**).

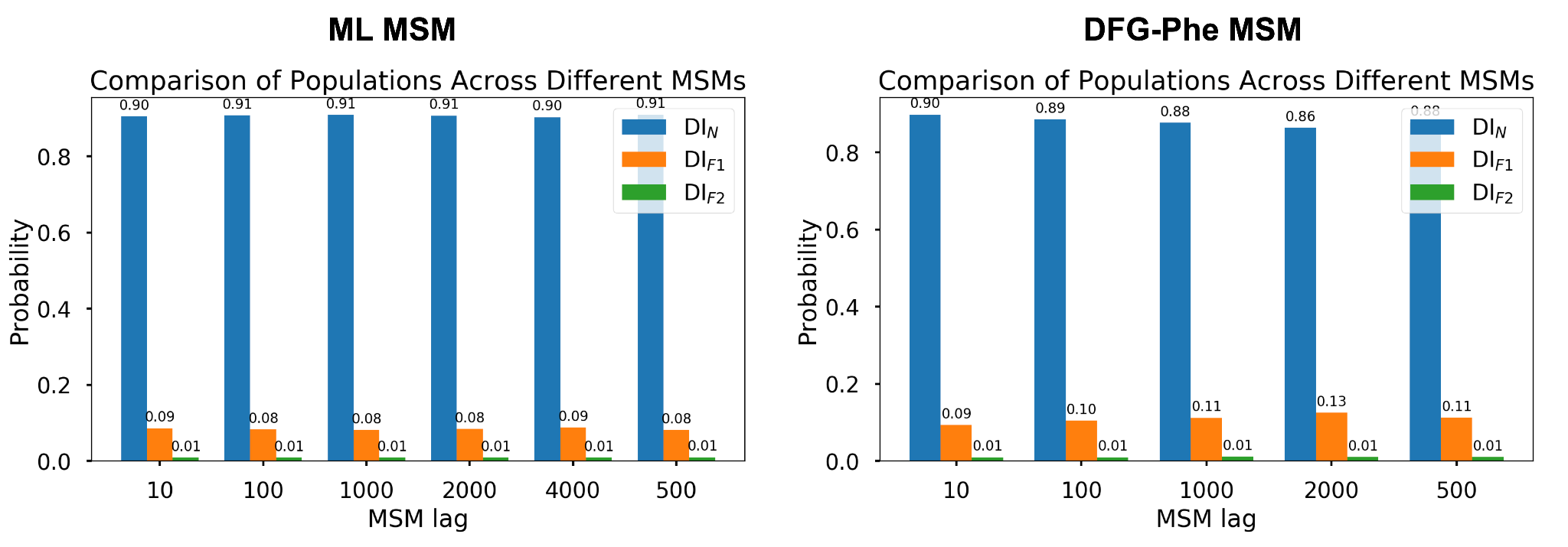

**Figure S9**. Analyzing the probability of DI_N/F1/F2_ conformational states as a function of MSM lag time (in frames) demonstrates the convergence of AI-augmented physics-based molecular simulations. MSM built on latent layers from ML model and sin/cos-transformed χ₁/χ₂ angles of DFG-Phe reveal similar relative populations of these conformational states.

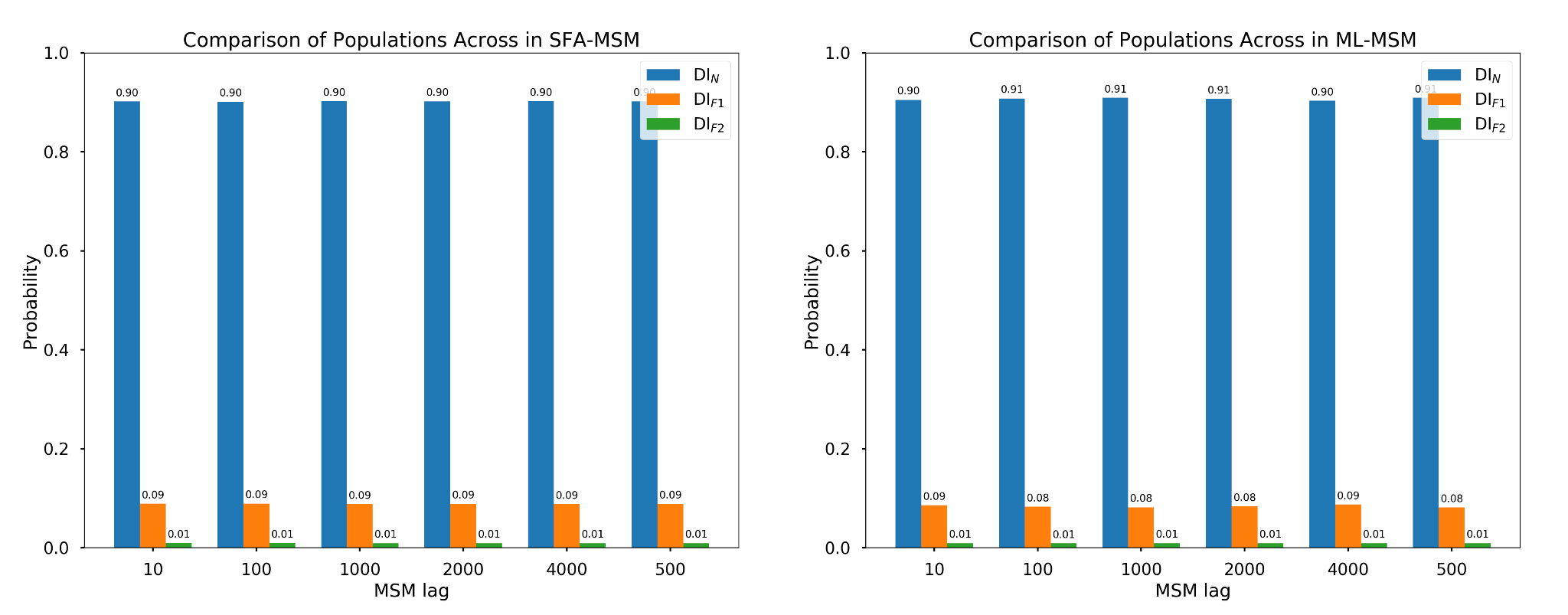

**Figure S10**. Analyzing the probability of DI_N/F1/F2_ conformational states as a function of MSM lag time (in frames) demonstrates the convergence of AI-augmented physics-based molecular simulations in apo RIPK2. MSM built on latent layers from ML model (ML-MSM) and first two slow features (SFA-MSM) reveal similar relative populations of these conformational states.

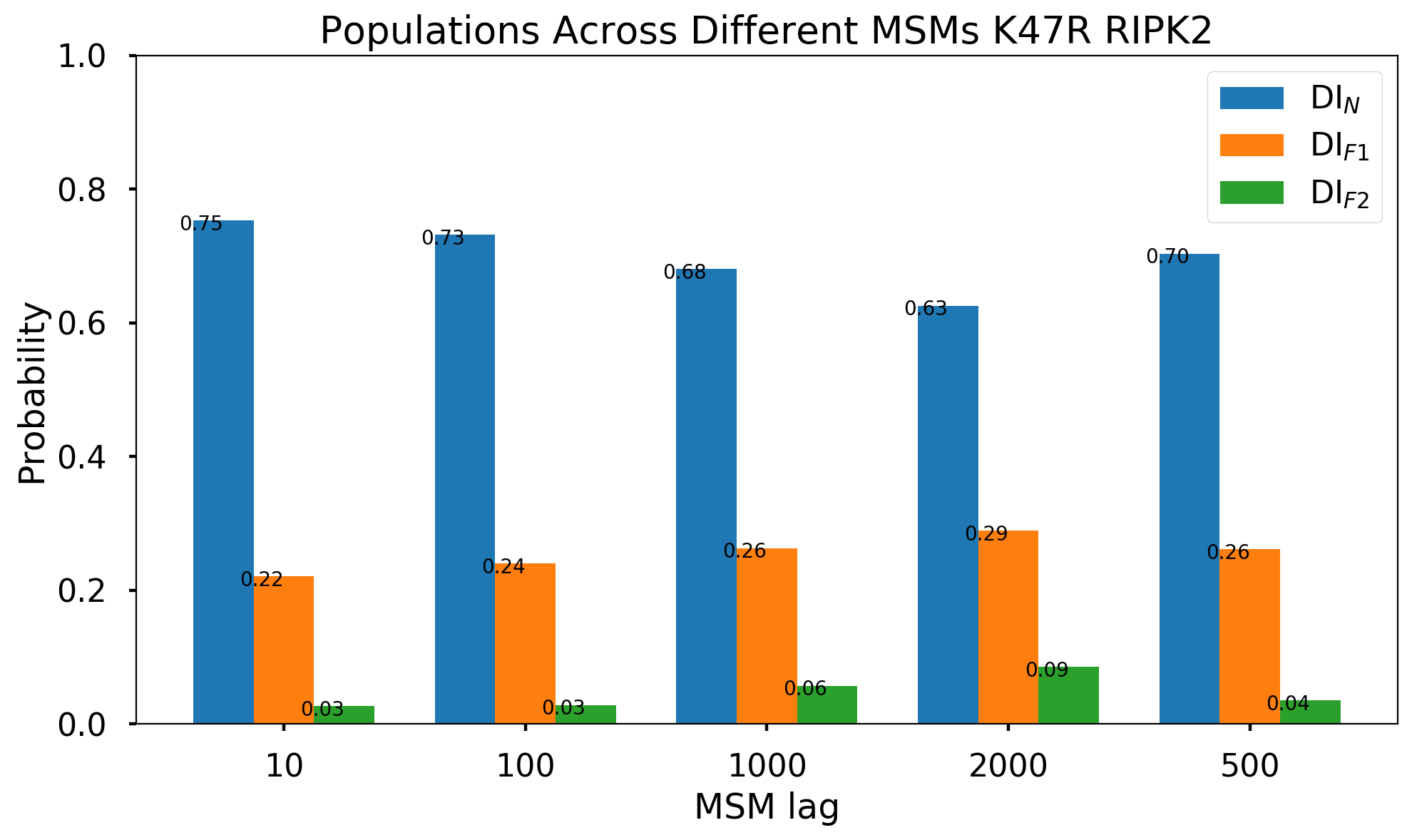

**Figure S11**. Probability of DI_N/F1/F2_ states as a function of lag time (in frames) for apo K47R RIPK2. The MSM was developed using sin/cos-transformed χ₁/χ₂ angles of DFG-Phe. Comparison with Figure S3 indicates that the K47R mutation shifts the population from DI_N_ to DI_F1/F2_.

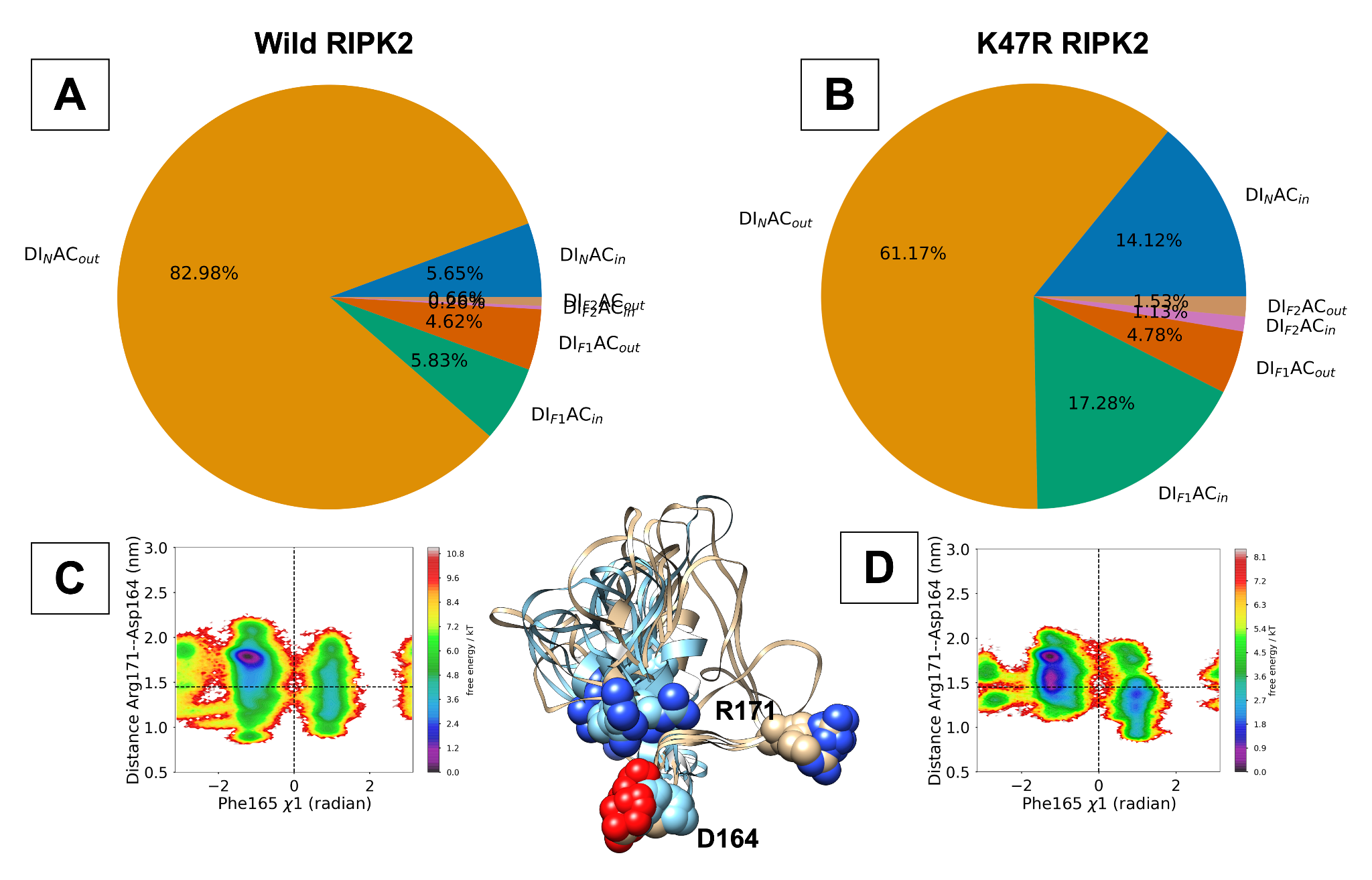

**Figure S12**. Population of AC_in/out_ conformations captured the *Cɑ--Cɑ* distance between Asp164 and Arg171 (silver: AC_out_, blue: AC_in_). MSM developed using χ₁/χ₂ angles of DFG-Phe projected on the Phe165 χ₁ angle and Arg171--Asp164 distance captures conformational heterogeneity. K47R mutation changes the conformation of the activation loop which shifts the population towards DI_F1_AC_in_ conformation. The crystal structure of K47R RIPK2 (PDB: 5NG3) captures a static structure of the DI_F1_AC_in_ ensemble.

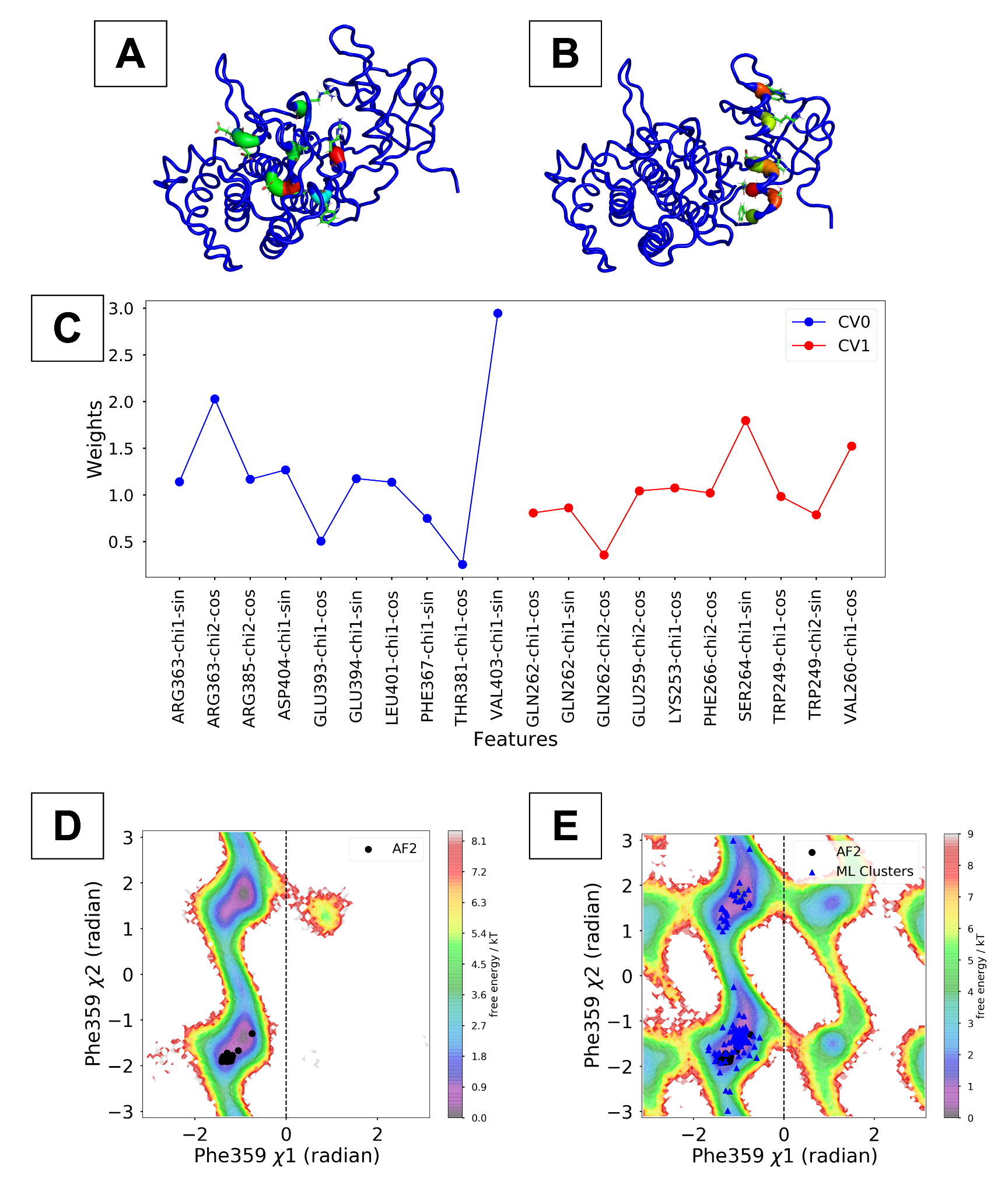

**Figure S13**. ML weights (C) projected onto two latent layers—CV0, which captures the DFG and activation loop dynamics, and CV1, which captures the α-C helix dynamics—reveal slowly varying structural features in 80 independent 10 ns MD simulations initiated from the AlphaFold predicted wild IRAK1 conformations. ML weights are mapped to the PDB structure (A, B) and highlighted as B-factors allowing for visualization of the key structural features captured by ML. Free energy surfaces projected along the χ₁ and χ₂ angles of Phe359 in IRAK1 compare the sampling efficiency of AlphaFold (AF)–seeded molecular simulations (D) with a machine-learning (ML)–augmented adaptive sampling (E) approach. Simulations initiated from the AF structure fail to capture the complete spectrum of DFG-Phe conformations in IRAK1. By contrast, the ML-augmented adaptive sampling launched from physics-refined conformational ensembles (highlighted by ML clusters in blue triangles) recovers the full range of conformational heterogeneity. Notably, the ML clusters encompass a broader spectrum of conformational diversity than the AlphaFold-based subsampling approach (black dots).

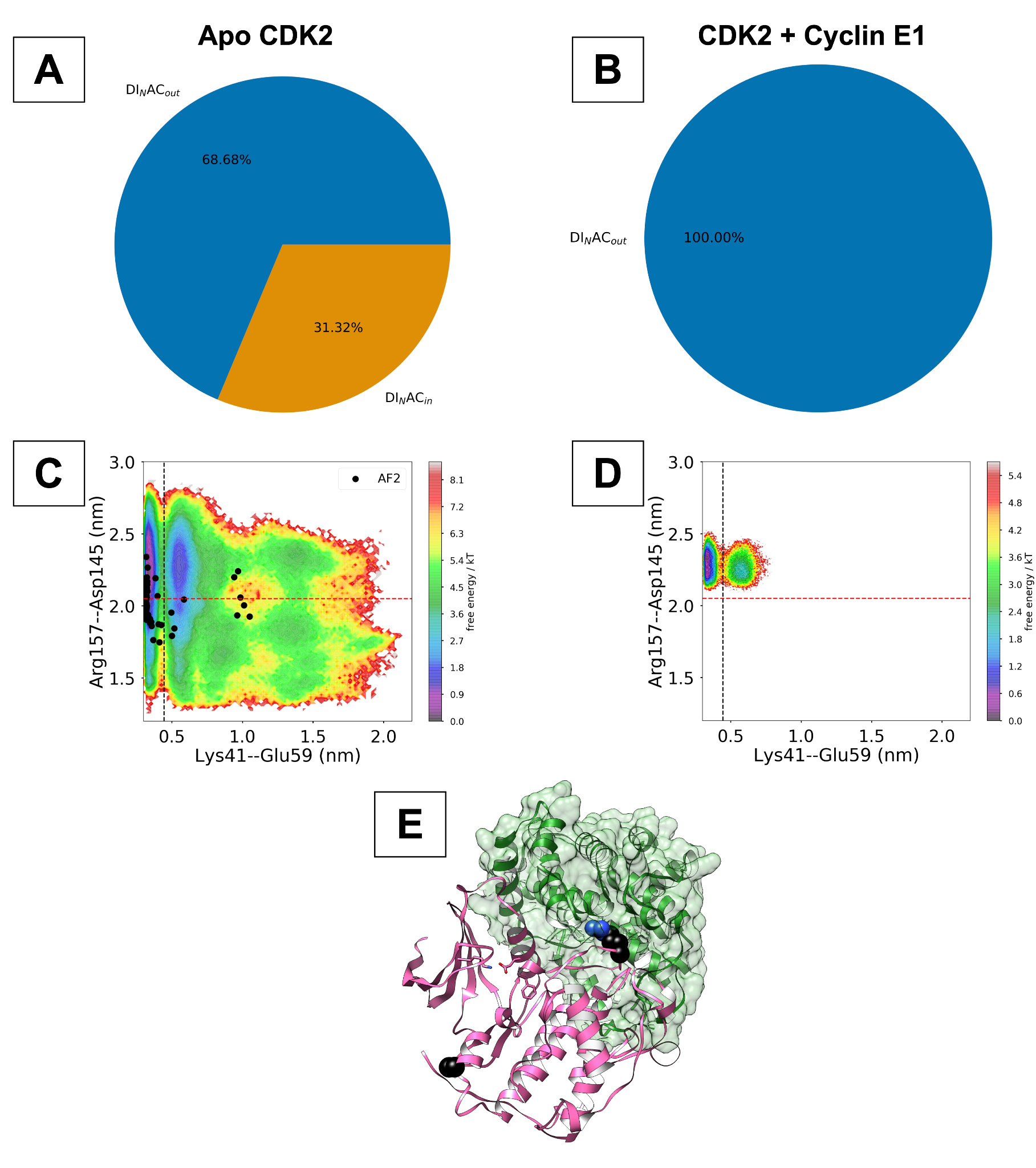

**Figure S14**. Physics based molecular simulation captures conformational heterogeneity of the activation loop in apo CDK2 and CDK2-CyclinE1 complex (A, B). The conformational dynamics of the activation loop is captured by the distance (Cɑ-Cɑ) between Asp145 and Arg157 (E). Molecular simulation also captures differentiable dynamics of Lys-Glu latched and unlatched conformations (C, D). Apo CDK2 remains in a dynamic equilibrium between AC_out_ and AC_in_ conformation (defined by the red dotted line which highlights a cut off of 3.5 nm). Binding of CyclinE1 stabilizes the AC_out_ conformation. The differentiable dynamics of the activation loop between CDKs and Cyclins will be consistent across the proteome.

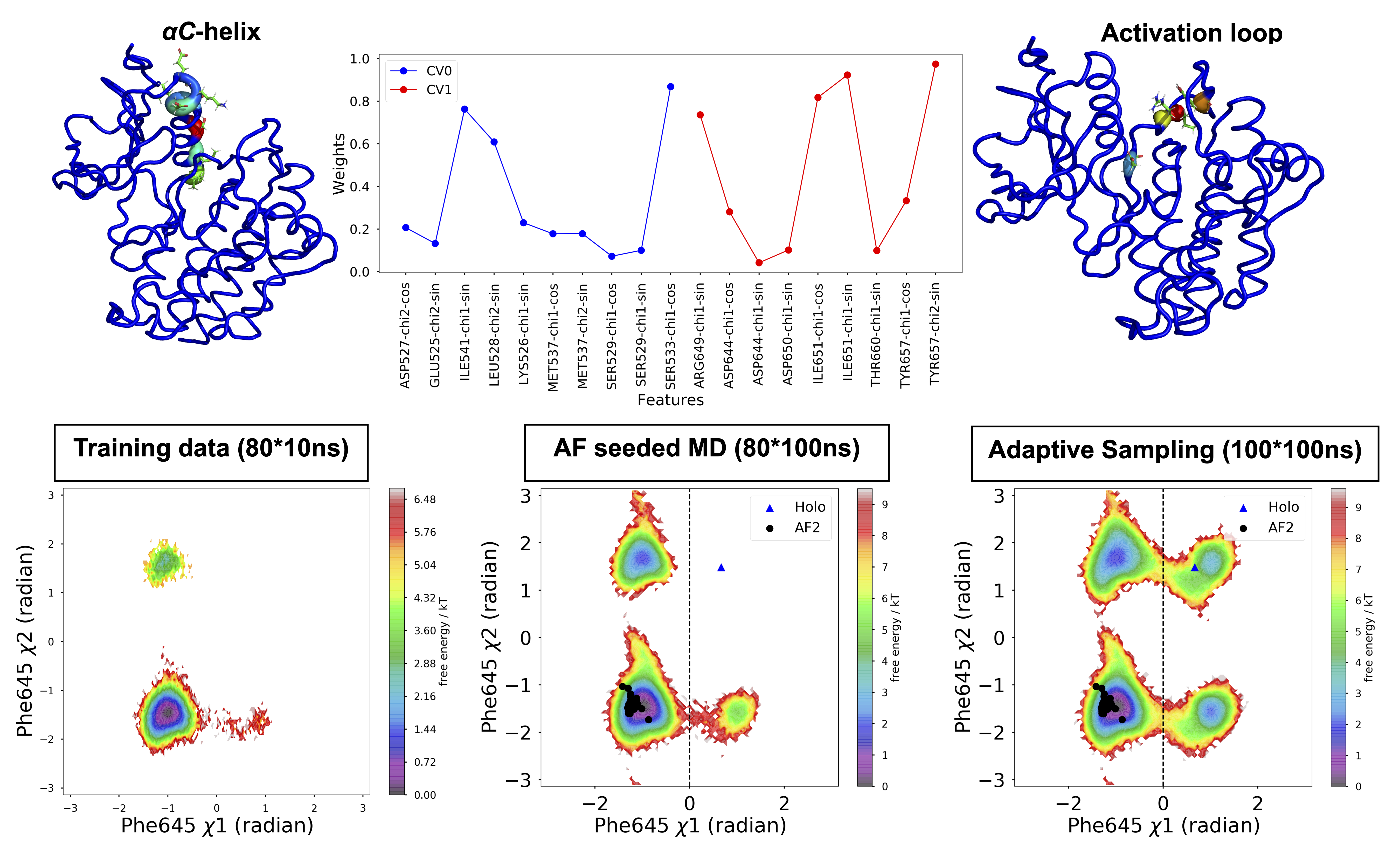

**Figure S15**. **Upper panel**: Time-lagged autoencoder trained on training data captures slowly varying structural features governing conformational dynamics of the *αC*-helix and the activation loop. ML weights projected on the protein structure highlights the key residues governing protein dynamics across different kinase domains. **Lower panel**: Projection of the free energy surface along the χ₁ and χ₂ angles of DFG-Phe in apo FGFR2 demonstrates how conformational space is sampled by the training data, AlphaFold (AF)-seeded MD simulations, and adaptive sampling approach. The training data were used to generate structures from the latent layer of the TAE, representing the physics-refined ensemble. Starting from this ensemble, the adaptive sampling protocol successfully captured dihedral flipping in FGFR2, including conformations stabilized by a selective FGFR2 inhibitor (holo). This outcome highlights the ability of our adaptive sampling strategy to explore the full spectrum of conformational space that AF-seeded MD simulations fail to sample.

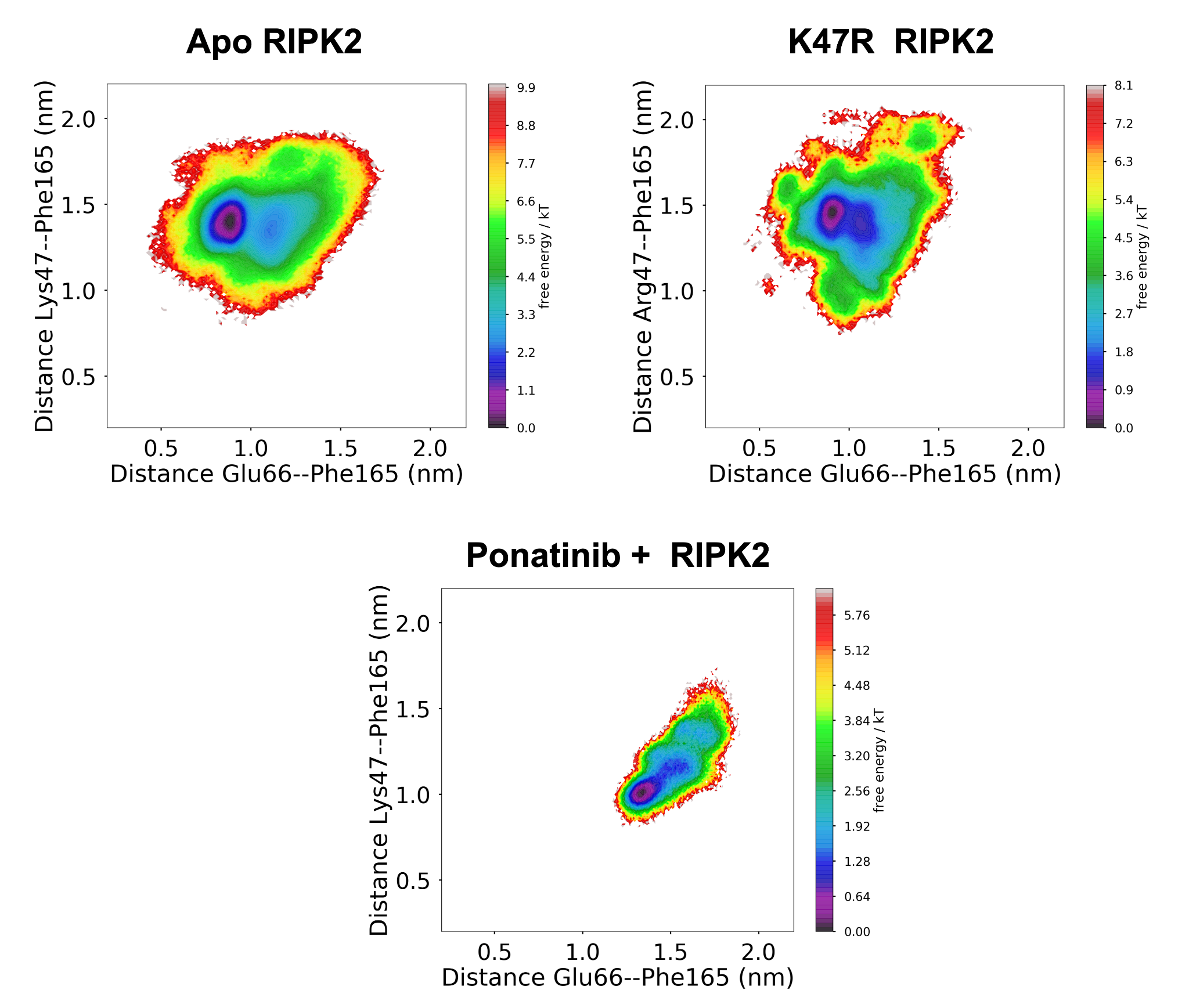

**Figure S16**. Conformational sampling of the DFG_in_ and DFG_out_ macrostates reveals that the apo and K47R mutant forms of RIPK2 stabilize the DI ensemble, whereas Ponatinib (a type II inhibitor)–bound RIPK2 stabilizes the DFG_out_ ensemble. However, classifying states solely as DFG_in_ and DFG_out_ does not fully capture the conformational heterogeneity within these ensembles. Figure S17 highlights how we can capture conformational heterogeneity within DFG_out_ ensemble.

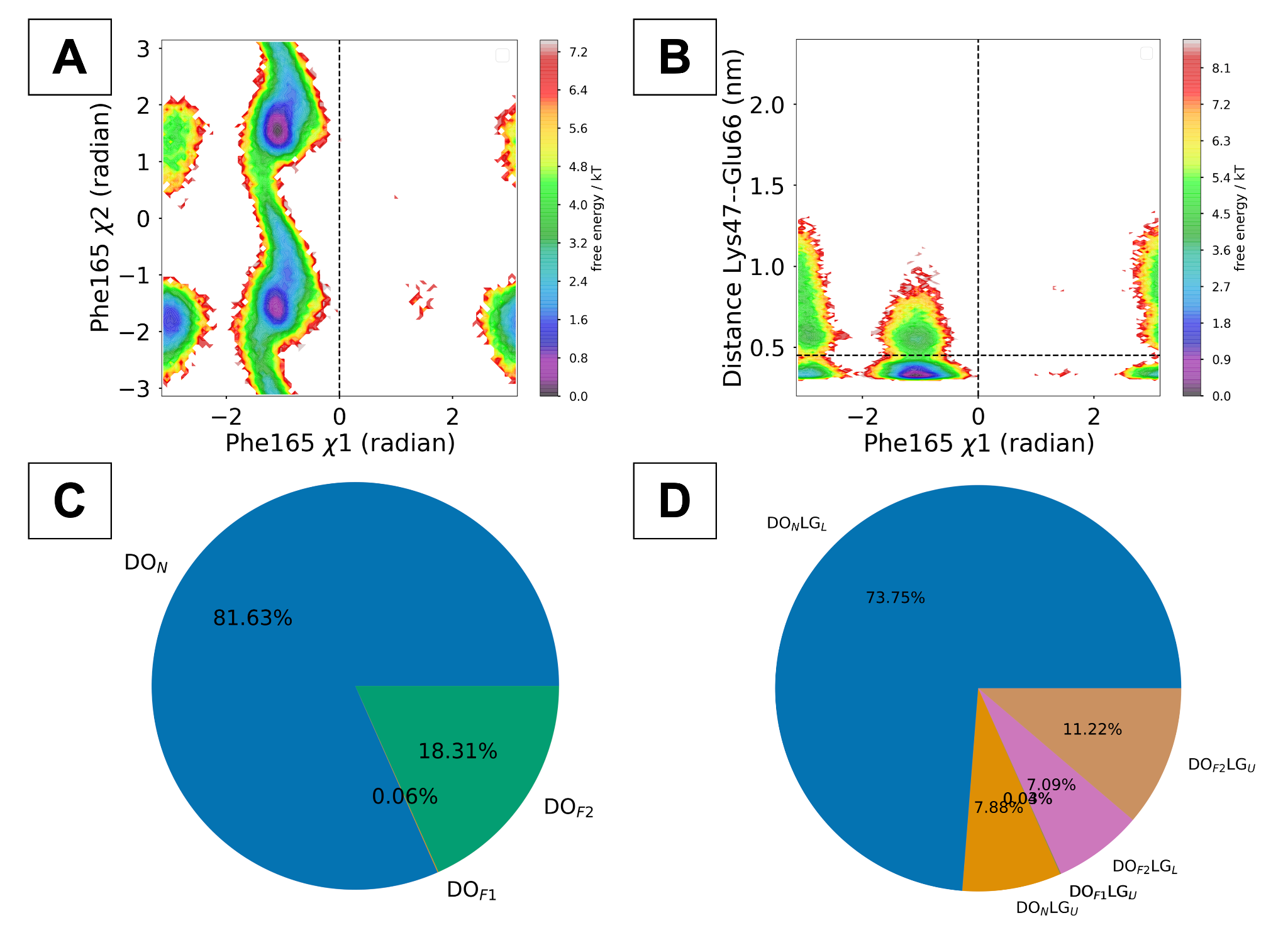

**Figure S17**. Physics-based molecular simulations of Ponatinib (type II inhibitor) bound RIPK2 reveal the conformational heterogeneity of the DFG-Phe in and highlight Lys–Glu (LG) interactions in DFG-out (DO) ensemble. These findings underscore the effectiveness of our ensemble definition in capturing conformational diversity within the DFG-out ensemble of kinases.

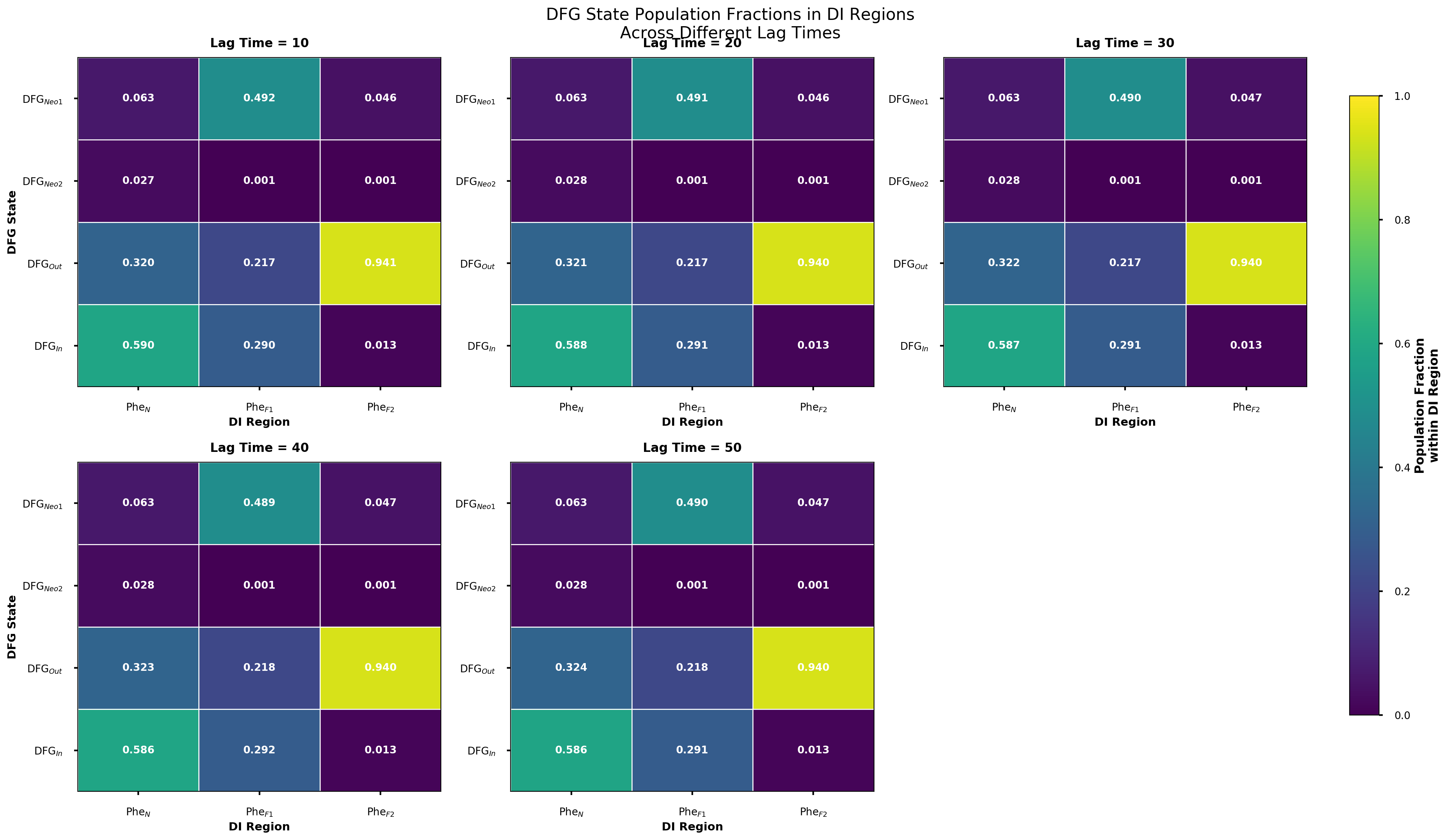

**Figure S18**. Heatmaps of the MSM‑weighted populations for the DFG_in_, DFG_out_, and DFG_Neo1/Neo2_ states—classified by the DFG‑Phe χ₁ dihedral—shows asymptotic values of populations across lag times, highlighting convergence of our simulation dataset. MSM was generated using sin and cos transformed χ₁ and χ_2_ angles of DFG-Phe.

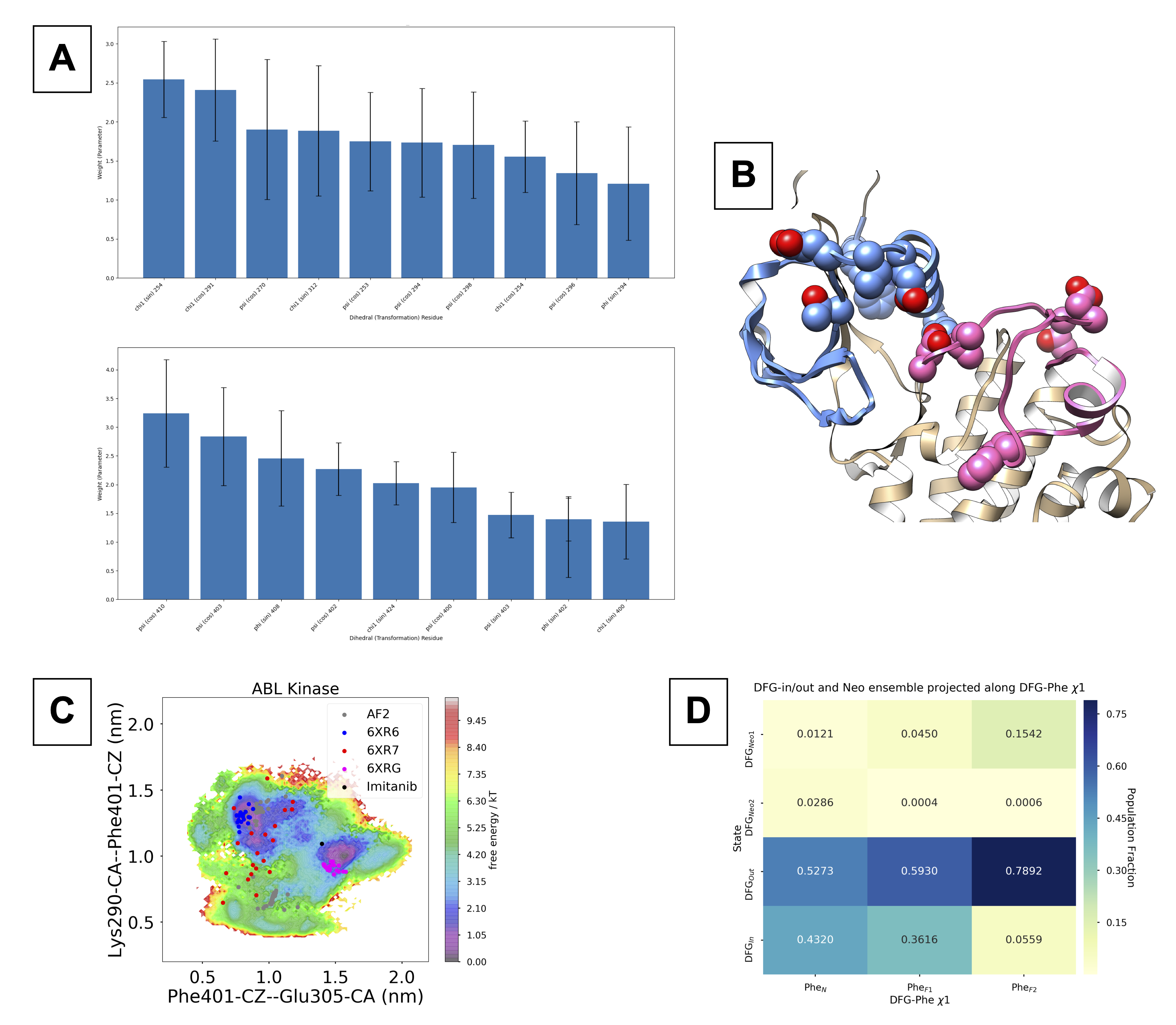

**Figure S19**. TAE weights of key residues (A) and the relative location of residues in different domains of ABL-kinase (B) highlighted by blue and magenta respectively. MSM trained on the TAE CVs and projected on the key distances highlighted conformational sampling of AlphaFold seeded molecular simulation (C). MSM population projected along DFG-substates (DFG_in/out/Neo1/Neo2_) vs DFG-Phe χ₁ angle (D) highlights the relative population of different conformational states within Phe_N/F1/F2_ ensemble (analogous to DI_N/F1/F2_).
